# Adipocyte Tribbles1 Regulates Plasma Adiponectin and Plasma Lipids in Mice

**DOI:** 10.1101/2021.03.11.434882

**Authors:** Elizabeth E. Ha, Gabriella I. Quartuccia, Ruifeng Ling, Chenyi Xue, Antonio Hernandez-Ono, Rami Imam, Jian Cui, Rajesh K. Soni, Robert C. Bauer

## Abstract

Multiple GWAS have identified SNPs in the 8q24 locus near the *TRIB1* gene that significantly associate with plasma lipids and coronary artery disease. While subsequent studies have uncovered roles for hepatic and myeloid *Trib1* in contributing to either plasma lipids or atherosclerosis, the causal tissue for these GWAS associations remains unclear. The same 8q24 SNPs significantly associate with plasma adiponectin levels in humans as well, suggesting a role for *TRIB1* in adipose tissue. Here, we report that adipocyte-specific *Trib1* knockout mice (Trib1_ASKO) have increased plasma adiponectin levels and decreased plasma cholesterol and triglycerides. We demonstrate that loss of *Trib1* increases adipocyte production and secretion of adiponectin independent of the known TRIB1 function of regulating proteasomal degradation. RNA-seq analysis of adipocytes and livers from Trib1_ASKO mice suggests that alterations in adipocyte function underlie the plasma lipid changes observed in these mice. Secretomics and RNA-seq analysis revealed that Trib1_ASKO mice have increased production of Lpl and decreased production of Angptl4 in adipose tissue, and fluorescent substrate assays confirm an increase in adipose tissue Lpl activity, which likely underlies the observed triglyceride phenotype. In summary, we demonstrate here a novel role for adipocyte *Trib1* in regulating plasma adiponectin, total cholesterol, and triglycerides in mice, confirming previous genetic associations observed in humans and providing a novel avenue through which *Trib1* regulates plasma lipids and coronary artery disease.

## Introduction

Plasma lipids, including triglycerides and cholesterol, are among the strongest risk factors for cardiovascular disease (CVD), and modulation of plasma lipid levels is among the most effective therapeutic strategies at combating atherosclerotic CVD. Multiple genome-wide association studies (GWAS) investigating cardiometabolic risk factors have identified SNPs in the 8q24 genomic locus that associate with plasma triglycerides (TGs), total cholesterol (TC), LDL-cholesterol (LDL-C), HDL-cholesterol (HDL-C), and coronary artery disease (CAD) [1-6], suggesting that this locus contains elements that regulate lipid metabolism and disease risk. These SNPs lie ∼40kb downstream of the Tribbles1 (*TRIB1*) gene, which codes for the TRIB1 pseudokinase. Studies in multiple genetic mouse models have since confirmed a role for both hepatic and macrophage *Trib1* in the regulation of lipid metabolism and CVD [7-9]. Viral mediated liver-specific overexpression of *Trib1* in mice was found to decrease plasma cholesterol and TGs, and hepatic deletion of *Trib1* increased plasma cholesterol and TG, while also causing hepatic steatosis due to increased *de novo* lipogenesis [7]. This latter phenotype confirmed an additional GWAS association between the 8q24 SNPs and circulating liver transaminases (ALTs/ASTs) [10], suggestive of a role for *TRIB1* in steatosis and hepatocellular health. A more recent study found that myeloid-specific *Trib1* knockout mice have reduced atherosclerotic burden [9] due to decreased OxLDL uptake by macrophages and reduced foam cell formation, highlighting the importance of tissue-specific gene functions as well as raising the question of possible roles for *Trib1* in other tissues in mediating the GWAS associations.

The same SNPs in the 8q24 locus that associate with plasma lipid traits, CAD, and ALTs have been found to additionally associate with plasma adiponectin levels in humans [11] (**Supplementary Figure 1**). Adiponectin is an adipokine, or signaling molecule secreted from adipocytes, that acts predominantly as an insulin sensitizing agent [12] but can also alter plasma lipids [13], hepatic fat content [14], and even CAD [15]. Given the 8q24 association and the fact that adiponectin is exclusively produced in adipocytes, we hypothesized that *TRIB1* plays a role in adipose tissue biology. Additionally, the myriad of roles for adiponectin in regulating cardiometabolic traits begs the question of whether the function of Trib1 in adipocytes is responsible for the observed metabolic genetic associations. Here, we report the first adipocyte-specific *Trib1* knockout mouse and show that these mice have increased plasma adiponectin levels and decreased plasma cholesterol and TG levels. Further mechanistic studies reveal that *Trib1* regulates plasma adiponectin through increased adiponectin production and secretion, and that *Trib1* modulates plasma TG clearance through regulation of adipose-specific lipoprotein lipase (Lpl) activity.

**Figure 1:**
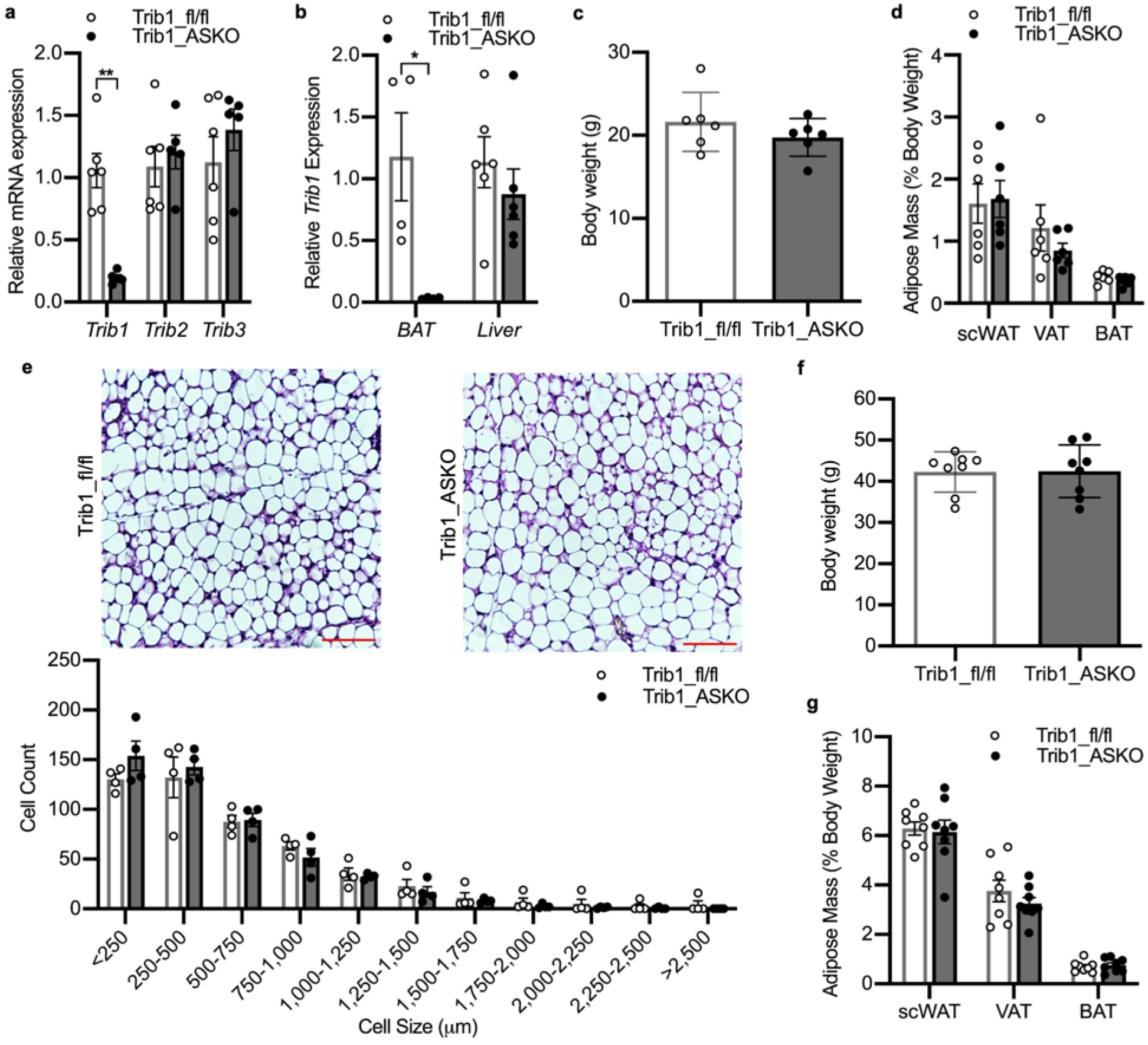
Adipocyte-specific knockout of Tribbles1 does not result in defects in adiposity. **a**, Taqman qPCR for *Trib1, Trib2, and Trib3* in scWAT from 8–10-week-old Trib1_fl/fl and ASKO mice (*n* = 5). **b**, Taqman qPCR for *Trib1* from BAT (*n* = 4) and livers (*n* = 6) of Trib1_fl/fl and Trib1_ASKO mice. **c**,**d**, Body weight (**c**) and adipose depot masses (**d**) in chow-fed Trib1_fl/fl and ASKO mice (*n* = 6). **e**, Representative H&E stain of scWAT from Trib1_fl/fl and ASKO mice and quantitation of cell size by Adiposoft (*n* = 4 mice). Bar = 100 μm. **f**,**g**, Body weight (**f**) and adipose depot masses (**g**) in 12 week HFD-fed Trib1_fl/fl and ASKO mice (*n* = 8). All gene expression data is depicted as mean ± s.e.m. All other data is depicted as mean ± s.d. Significance in all panels determined by Student’ s *t* test (*p <0.05, **p < 0.01).

## Results

### Adipocyte-specific Trib1 knockout does not alter body weight, adiposity, or adipose inflammation

We generated *Trib1* adipocyte-specific knockout (Trib1_ASKO) mice by crossing previously described Trib1-floxed (Trib1_fl/fl) C57BL/6 mice [7] with transgenic mice expressing Cre recombinase under the adipocyte-specific *Adipoq* promoter. Efficient *Trib1* deletion in adipose tissue of Trib1_ASKO mice was confirmed by qPCR in both subcutaneous white adipose tissue (scWAT) (**Figure 1a**) and brown adipose tissue (BAT) (**Figure 1b**), and we did not detect any compensatory changes in *Trib2* or *Trib3* expression in scWAT (**Figure 1a**). Trib1 message was unchanged in other tissues, including the livers (**Figure 1b**) of Trib1_ASKO mice, confirming specificity of the model. Chow-fed Trib1_ASKO mice had similar overall body weight and fat pad mass to Trib1_fl/fl mice (**Figure 1c,d**), and there was no difference in adipocyte morphology or size as measured by H&E staining and subsequent morphometric analysis (**Figure 1e**). Similar results were observed in mice fed a 45% kcal high-fat diet (HFD) for 12 weeks (**Figure 1f,g**).

To better understand the effects of *Adipoq-*Cre mediated knockout of *Trib1* in adipocytes, we utilized an *in vitro* model of adipocyte culture, where we isolated the stromal vascular fraction (SVF) from the scWAT of Trib1_fl/fl and Trib1_ASKO mice and differentiated them to adipocytes. Consistent with the lack of an adiposity phenotype in the adult mice, SVF-derived adipocytes from Trib1_fl/fl and ASKO mice differentiated similarly, as assessed by time course gene expression of adipogenic markers *Pparg, Cebpa*, and *Adipoq*, as well as by cellular morphology and lipid accumulation (**Supplemental Figure 2**). Importantly, *Trib1* expression in adipocytes derived from ASKO mice was lower relative to adipocytes derived from Trib1_fl/fl mice (**Supplemental Figure 2a**), demonstrating that *Adipoq*-Cre is efficiently expressed in the *in vitro* setting upon differentiation.

**Figure 2:**
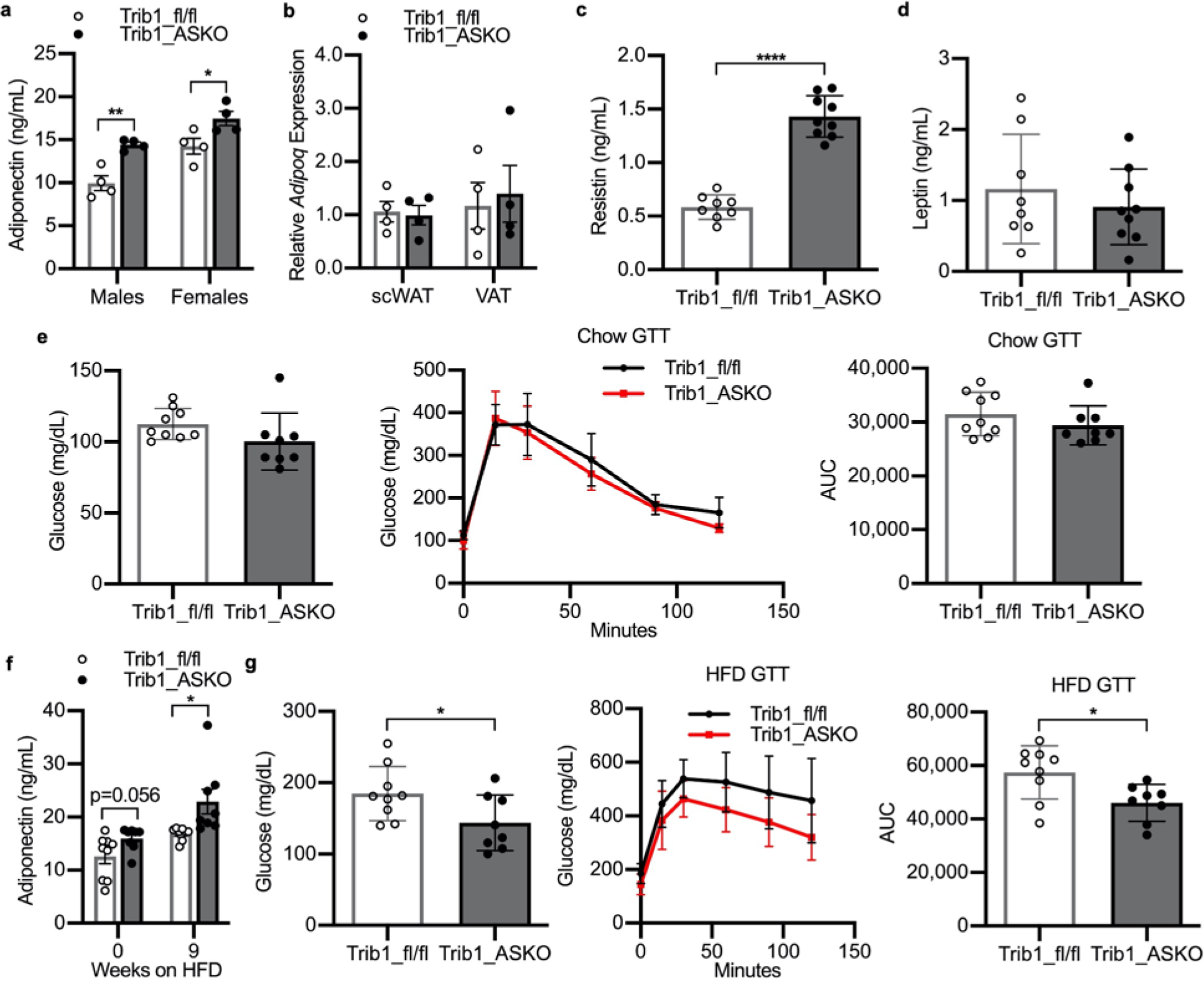
Trib1_ASKO mice have increased plasma adiponectin. **a**, Plasma adiponectin in 4hr-fasted 8-week-old chow-fed Trib1_fl/fl and Trib1_ASKO mice (*n* = 4). **b**, Taqman qPCR for *Adipoq* in scWAT and VAT from Trib1_fl/fl and Trib1_ASKO mice (*n* = 4). **c**,**d**, Plasma resistin and plasma leptin (**d**) in 4 hr-fasted chow-fed Trib1_fl/fl and Trib1_ASKO mice (*n* = 8). **e**, Glycemic traits after 16 hr overnight fast in chow-fed 8–10-week-old male Trib1_fl/fl and Trib1_ASKO mice (*n* = 7). **f**, Plasma adiponectin in 4 hr-fasted 9 week HFD-fed Trib1_fl/fl and Trib1_ASKO mice (*n* = 8). **g**, Glycemic traits after 16 hr overnight fast in 12 week HFD-fed male Trib1_fl/fl and Trib1_ASKO mice (*n* = 8). Gene expression is depicted as mean ± s.e.m. All other data is depicted as mean ± s.d. Significance in all panels determined by Student’ s *t* test (*p <0.05, **p < 0.01, ****p < 0.0001).

A previous study reported that Trib1 haploinsufficiency in mice impairs the upregulation of inflammatory genes in adipose in response to proinflammatory stimuli such as LPS, TNF-α, and high-fat diet feeding [16]. Given the known contribution of adipose tissue inflammation to obesity and metabolic disease, we checked if Trib1_ASKO mice had decreased inflammatory markers in adipose. We measured inflammatory gene expression in adipose tissue in both chow-fed and HFD-fed conditions and observed no difference between the groups in either diet setting (**Supplemental Figure 3a,b**). Similarly, we did not observe any changes in the transcriptional response to TNF-α treatment in SVF-derived adipocytes from Trib1_ASKO mice compared to Trib1_fl/fl controls (**Supplemental Figure 3c**). These data suggest that the phenotypes we have observed in our mice are not due to changes in adipose inflammation.

**Figure 3:**
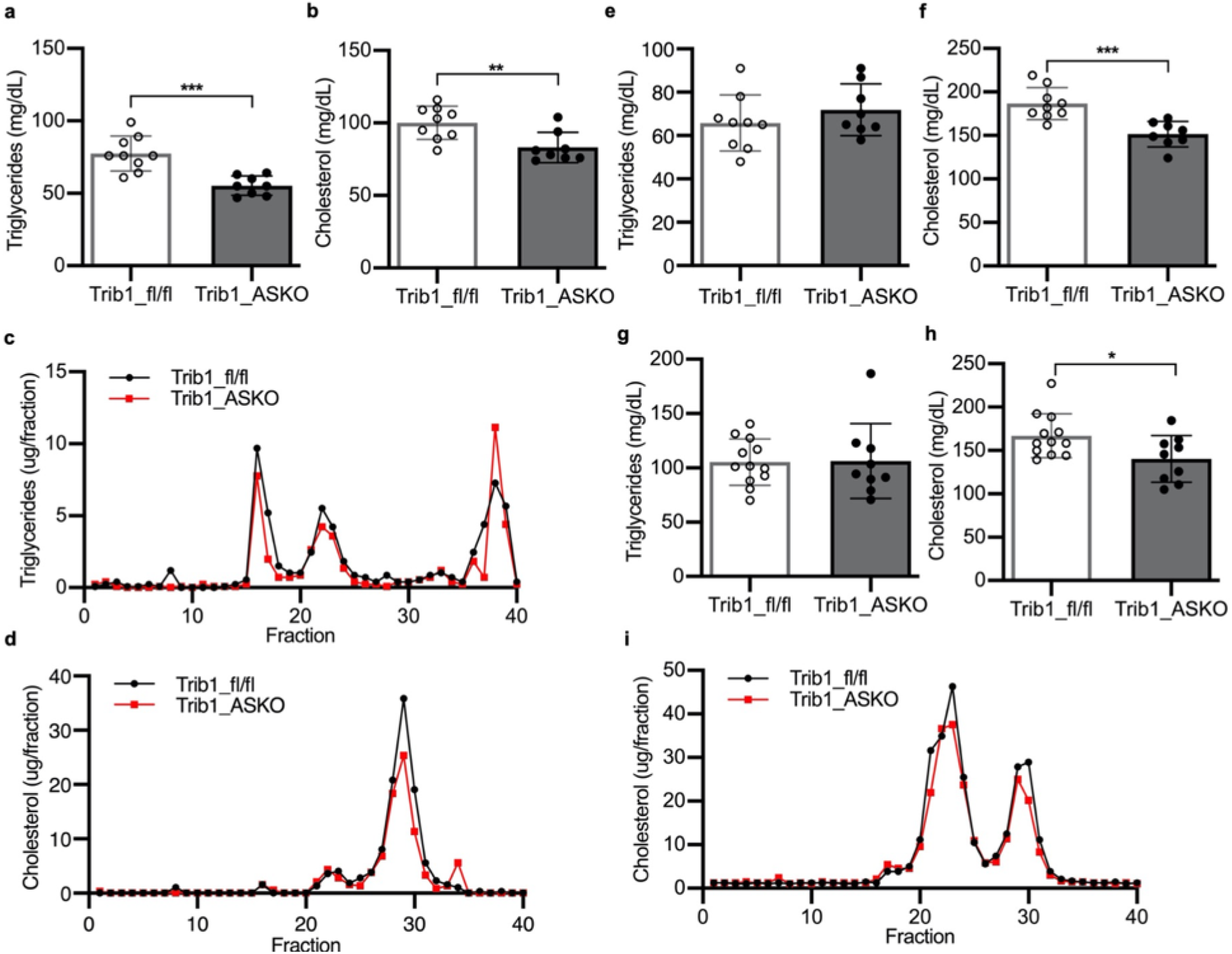
Trib1_ASKO mice have decreased plasma cholesterol and triglycerides. **a**,**b**, Plasma triglyceride (**a**) and total cholesterol (**b**) levels in 8–10-week-old, 4 hr-fasted chow-fed male Trib1_fl/fl and Trib1_ASKO mice (*n* = 8). **c**,**d**, Plasma triglyceride (**c**) and cholesterol (**d**) FPLC profiles of pooled plasma (*n* = 4) from 4 hr-fasted chow-fed male Trib1_fl/fl and Trib1_ASKO mice. **e**,**f**, Plasma triglyceride (**e**) and total cholesterol (**f**) levels in 4 hr-fasted 12 week HFD-fed Trib1_fl/fl and Trib1_ASKO mice (*n* = 8). **g**,**h**, Plasma triglyceride(**g**) and total cholesterol (**h**) levels in 8-week-old, 4 hr-fasted chow-fed male Trib1_fl/fl; Ldlr KO and Trib1_ASKO; Ldlr KO mice (*n =* 9). **i**, Cholesterol FPLC profile of pooled plasma (*n* = 4) from 4 hr-fasted chow-fed female Trib1_fl/fl; Ldlr KO and Trib1_ASKO; Ldlr KO mice. Data is depicted as mean ± s.d. Significance in all panels determined by Student’ s *t* test (*p < 0.05, **\*\***p < 0.01, *** p < 0.001).

### Adipocyte-specific Trib1 knockout mice have increased plasma adiponectin and decreased plasma lipids

Given the association in humans between SNPs near *TRIB1* and plasma adiponectin, we first sought to determine if Trib1_ASKO mice had altered circulating adiponectin levels. We found that both male and female Trib1_ASKO mice on chow diet have significantly increased plasma adiponectin (>20%) levels compared to wild-type counterparts (**Figure 2a**). The increase in plasma adiponectin was not accompanied by detectable changes in adiponectin message levels in the scWAT or visceral adipose tissue (VAT) (**Figure 2b**), suggesting a posttranscriptional role for Trib1 in plasma adiponectin regulation. We checked the plasma levels of other adipokines and found that plasma resistin levels were also increased in Trib1_ASKO mice (**Figure 2c**). However, plasma levels of leptin, another abundant adipokine, were not significantly changed in Trib1_ASKO mice (**Figure 2d**), demonstrating that Trib1 regulates the secretion of specific adipokines and not global adipokine secretion. Despite increased adiponectin levels, glucose tolerance was not significantly changed in 8–12-week-old chow-fed mice (**Figure 2e**), and SVF-derived adipocytes did not demonstrate increased insulin signaling upon insulin stimulation (**Supplemental Figure 4a**). HFD-fed Trib1_ASKO mice maintained increased adiponectin levels (**Figure 2f**), and, in contrast to chow-fed mice, these mice also had significantly improved glucose tolerance (**Figure 2g**) as well as decreased fasting plasma insulin levels (**Supplemental Figure 4b**), consistent with studies that show association between increased plasma adiponectin levels and improved insulin sensitivity [12].

**Figure 4:**
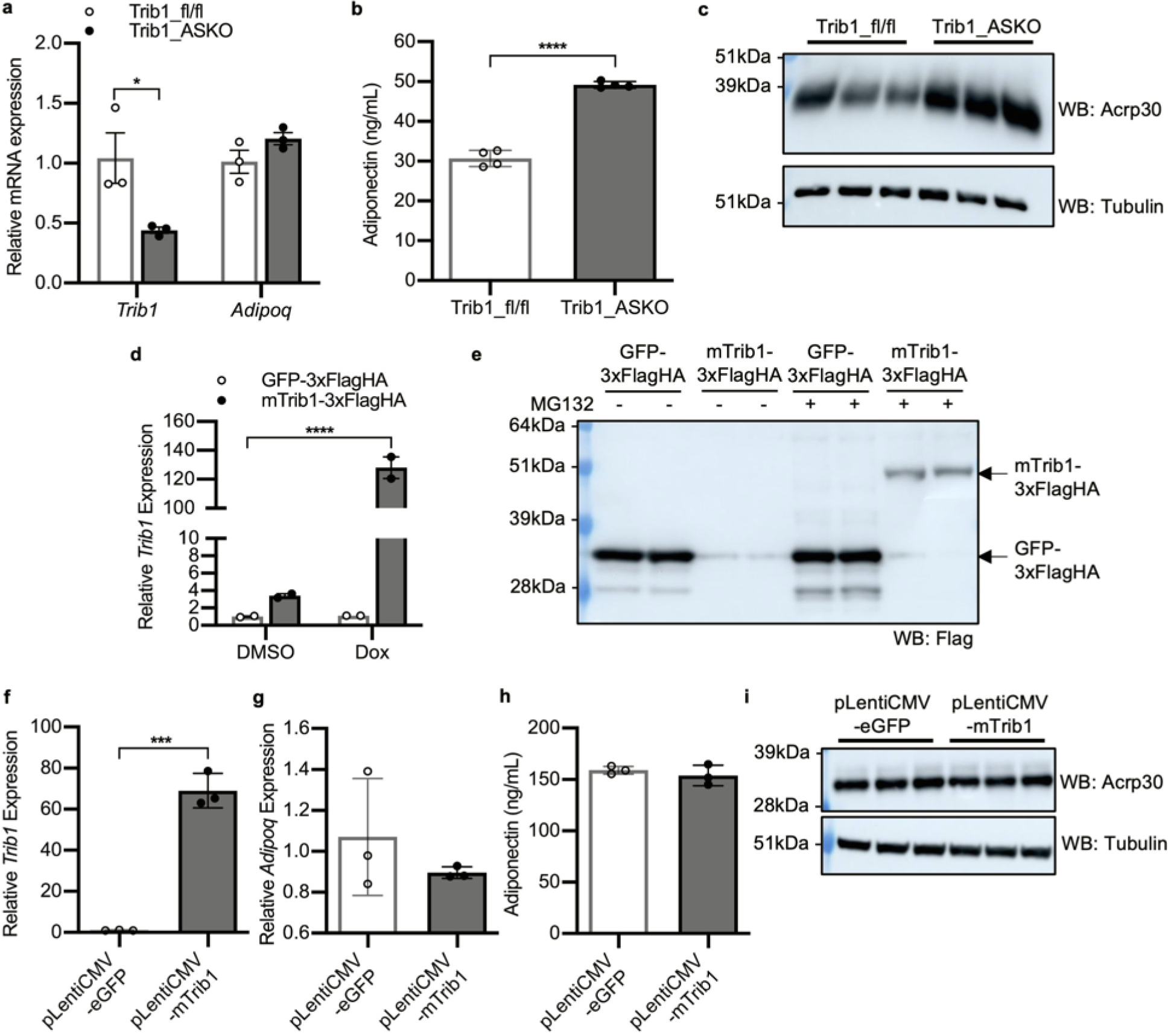
Adipocyte-specific knockout of Tribbles1 results in increased adiponectin secretion. **a**, Taqman qPCR for *Trib1* and *Adipoq* in SVF-derived adipocytes (*n* = 3). **b**, Adiponectin concentration in conditioned media from SVF-derived adipocytes (*n* = 4). Conditioned media was generated by culturing SVF-derived adipocytes in OptiMEM reduced-serum media for 4 hr. **c**, Western blot analysis of adiponectin (Acrp30) and tubulin levels in SVF-derived adipocytes. **d**, Taqman qPCR for *Trib1* in pSlik-neo-TTMCS_eGFP-3xFlagHA and pSlik-neo-TTMCS_mTrib1-3xFlagHA stable 3T3-L1 cells treated with either DMSO or doxycycline (1 μg/mL) (*n* = 2). Gene expression is expressed relative to the DMSO-treated GFP stable cells. Significance relative to DMSO treated GFP stable cells was determined by 1-way ANOVA (Dunnett’ s multiple comparison test) **e**, Western blot for Flag-tagged protein overexpression in pSlik-neo-TTMCS_eGFP-3xFlagHA and pSlik-neo-TTMCS_mTrib1-3xFlagHA stable 3T3-L1 preadipocytes induced with 1 μg/mL Dox for 48 hr and treated with or without 20 μM MG132 for 5 hr. **f-i**, Mature 3T3-L1 adipocytes were transduced with lentivirus to overexpress eGFP (pLentiCMV-eGFP) or mTrib1 (pLentiCMV-mTrib1) under the CMV promoter. Taqman qPCR for *Trib1* (**f**) and *Adipoq* (**g**) (*n* = 3). Gene expression is expressed relative to the pLentiCMV-eGFP group. **h**, ELISA for adiponectin in 4 hr conditioned media (*n* = 3). **i**, Western blot for adiponectin protein expression. Gene expression is depicted as mean ± s.e.m. All other data is depicted as mean ± s.d. Significance in all panels determined by Student’ s *t* test except where indicated (*p<0.05, ***p<0.001, ****p<0.0001).

Since the same SNPs in the 8q24 locus significantly associate with both plasma adiponectin and plasma lipids (LDL-C, HDL-C, and TG), we next asked if adipocyte *Trib1* contributes to plasma lipid regulation. We found that chow-fed Trib1_ASKO mice display decreased plasma TG (>28%) and TC (15%) levels compared to wild-type counterparts (**Figure 3a,b**), demonstrating a role for adipocyte *Trib1* in plasma lipid regulation. We note that this phenotype of decreased plasma lipids is the opposite direction of the effect of the liver-specific knockout of *Trib1*, which results in increased plasma lipids [7], demonstrating opposing tissue-specific roles for Trib1 in regulating plasma lipids. FPLC analysis of pooled plasma revealed that Trib1_ASKO mice have reduced cholesterol in the HDL fraction as well as decreased TGs in both the VLDL and LDL fractions (**Figure 3c,d**). Trib1_ASKO mice continue to demonstrate lower total plasma cholesterol when placed on HFD for 12 weeks, although TGs normalized to WT levels (**Figure 3e,f**). To test for involvement of the LDL receptor in the cholesterol phenotype, we crossed the Trib1_ASKO mice to Ldlr KO mice. While plasma TGs did not differ between the groups (**Figure 3g**), we found that Trib1_ASKO Ldlr KO mice on chow diet had significantly decreased total cholesterol compared to Trib1_fl/fl Ldlr KO mice (**Figure 3h**), demonstrating that the cholesterol phenotype is at least partially independent of the LDL receptor pathway. FPLC analysis of pooled plasma from Trib1_ASKO; Ldlr KO mice on chow diet further revealed decreased cholesterol levels in both the LDL and HDL fractions in ASKO mice (**Figure 3i**).

### Trib1 in adipocytes regulates adiponectin secretion in a posttranscriptional and proteasome-independent mechanism

Since plasma adiponectin levels were increased in Trib1_ASKO mice, we hypothesized that *Trib1* deficiency in adipocytes promotes increased adiponectin secretion. To test this hypothesis, we investigated adiponectin protein expression and secretion from SVF-derived adipocytes from Trib1_fl/fl and Trib1_ASKO scWAT. Consistent with observations in whole adipose tissue, adiponectin mRNA expression was unchanged in SVF-derived adipocytes (**Figure 4a**). However, adiponectin was increased in the conditioned media above adipocytes derived from Trib1_ASKO SVF compared to Trib1_fl/fl SVF (**Figure 4b**), confirming increased adiponectin secretion. This was also accompanied by a clear increase in intracellular adiponectin protein levels (**Figure 4c**), suggesting that increased secretion is in part due to increased cellular adiponectin protein levels, despite the lack of a transcriptional change.

We next asked whether *Trib1* overexpression in adipocytes would also have an effect on adiponectin secretion and intracellular protein. We generated a doxycycline-inducible 3xFlagHA-tagged *Trib1* overexpression 3T3-L1 stable cell line that was able to overexpress *Trib1* >100-fold over wild-type values (**Figure 4d)**. We found that Trib1 protein was not detectable in these cells via western blot unless the cells were first treated with the proteasome inhibitor MG132, suggesting that Trib1 is unstable and undergoes rapid proteasomal degradation (**Figure 4e**) in 3T3-L1 cells. To avoid differences in cell line differentiation capacity caused by selection, we ultimately used lentiviral delivery of *Trib1* and e*GFP* expressed under the CMV promoter in mature 3T3-L1 adipocytes to assess the effects of *Trib1* overexpression in culture. We achieved >60-fold overexpression of *Trib1* via this method, but observed no changes in adiponectin secretion or protein levels compared to the GFP control (**Figure 4f-i**). Thus, while we were able to show that *Trib1* deficiency robustly affects adiponectin protein and secretion in adipocytes, we were unable to produce any effect on adiponectin with *Trib1* overexpression *in vitro*.

To better understand the molecular function of Trib1 in adipose tissue and how it may be regulating adiponectin, we first investigated previously reported functions of Trib1 described in other models. As a pseudokinase, Trib1 lacks catalytic phosphorylation activity, and is instead understood to function as a scaffolding protein that mediates interactions between its binding partners [17]. In this regard, Trib1 is best known for its role in the proteasomal degradation of the transcription factor C/EBPα via mediating its ubiquitination by the COP1 E3 ubiquitin ligase [18]. In keeping with that function, we found that C/EBPα protein levels were increased in the adipose tissue of Trib1_ASKO mice (**Figure 5a**) without observable changes in *Cebpa* message levels (**Figure 5b, Supplemental Figure 2c**). Although C/EBPα is a known transcriptional regulator of adiponectin expression [19], we did not observe a consistent increase in adiponectin expression in either tissue or SVF-derived adipocytes (**Figure 2b, Figure 4a**), consistent with *Trib1* regulating adiponectin through a mechanism independent of C/EBPα-mediated transcription.

**Figure 5:**
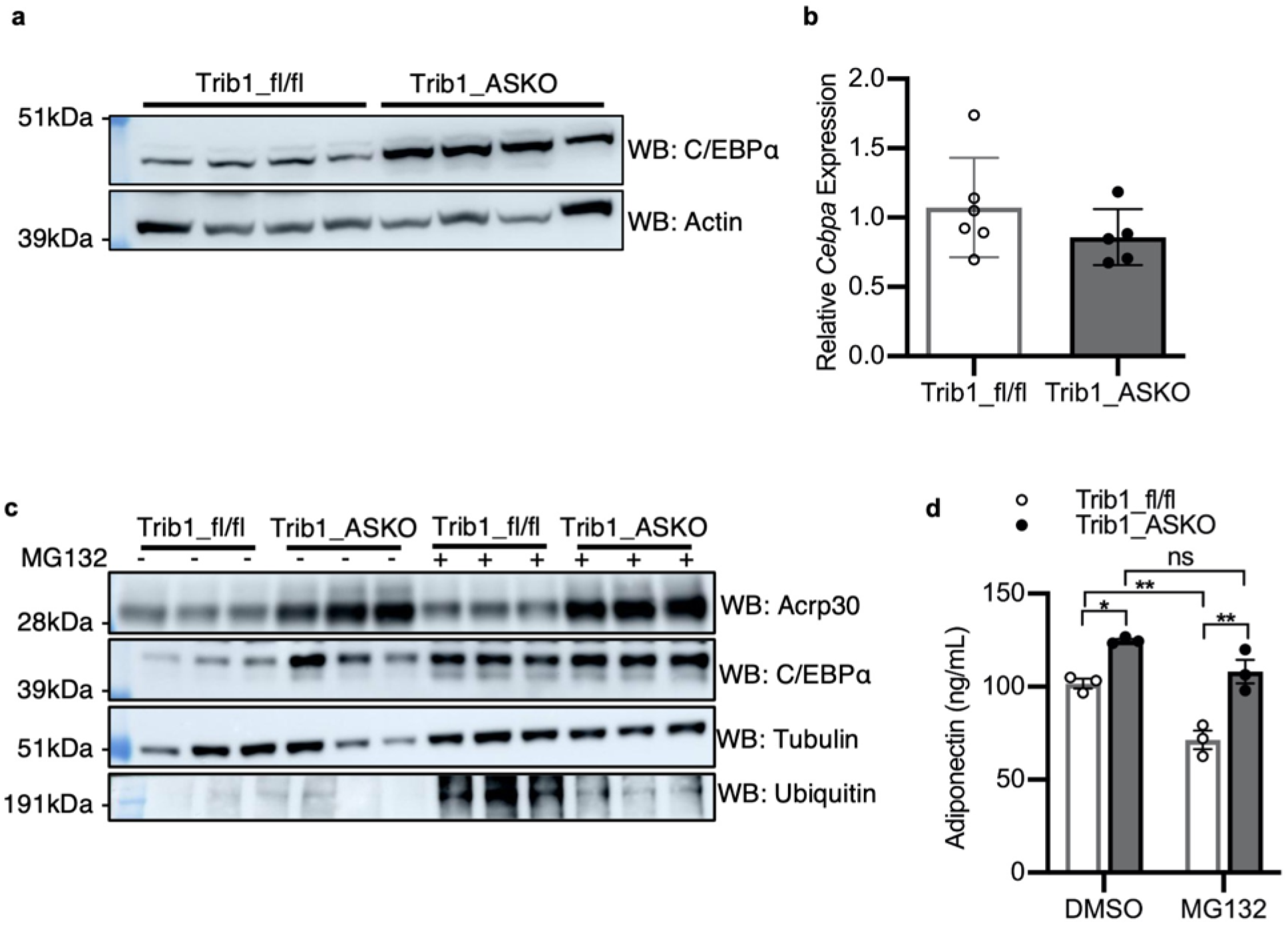
Trib1 does not regulate adiponectin through the proteasome in SVF-derived adipocytes. **a**, Western blot of C/EBPα in scWAT of Trib1_fl/fl and Trib1_ASKO mice. **b**, qPCR for *Cebpa* gene expression in scWAT of Trib1_fl/fl and Trib1_ASKO mice (*n* = 5). **c**,**d**, SVF-derived adipocytes from Trib1_fl/fl and Trib1_ASKO scWAT were differentiated, pretreated with 30 μM MG132 for 1hr, and then treated with 30 μM MG132 for an additional 4 hr before measuring protein expression and adiponectin secretion. **c**, Western blot for adiponectin (Acrp30) and C/EBPα protein in 5 hr MG132 treated Trib1_fl/fl and Trib1_ASKO adipocytes. **d**, ELISA for adiponectin in 4 hr conditioned media from 30 μM MG132 treated Trib1_fl/fl and Trib1_ASKO adipocytes (*n* = 3). Gene expression is depicted as mean ± s.e.m. All other data is depicted as mean ± s.d. Significance in (**b**) determined by Student’ s *t* test, and significance in by 2-way ANOVA (Tukey’ s multiple correction) (ns = not significant, *p < 0.05, **p < 0.01).

Given Trib1’ s role in mediating ubiquitination of proteins for proteasomal degradation, we further asked if the proteasome was important in Trib1’ s regulation of C/EBPα and adiponectin. We treated Trib1_ASKO and Trib1_fl/fl SVF-derived adipocytes with MG132 to determine if proteasomal inhibition would increase C/EBPα and adiponectin protein levels in the control cells but not the KO cells, normalizing the protein levels between the two. We found that MG132 treatment did normalize C/EBPα protein levels (**Figure 5c**) between the two groups, consistent with the known function of Trib1 regulating C/EBPα degradation. However, MG132 treatment did not affect the difference in adiponectin secretion (**Figure 5d)** or protein levels (**Figure 5c**) between control and ASKO cells, suggesting that *Trib1* is regulating adiponectin through a proteasome and C/EBPα-independent pathway.

### RNA-sequencing of adipocytes and hepatocytes reveals a primary role for adipose tissue in altered plasma lipid metabolism in Trib1_ASKO mice

To understand the mechanism by which adipocyte-specific *Trib1* regulates plasma lipids, we sequenced RNA from adipocytes isolated from the scWAT of Trib1_fl/fl and Trib1_ASKO mice. Differential expression analysis revealed over 2000 genes that were differentially expressed at a greater than 2-fold change (**Figure 6a**), emphasizing a widespread role for Trib1 in adipose. We considered the possibility that altered hepatic metabolism could explain the phenotypes observed in Trib1_ASKO mice, given that the liver is a major regulator of lipoprotein metabolism and that adipokines such as adiponectin can signal to the liver. However, RNA-seq of livers from the same mice revealed very few differentially expressed genes, none of which were major lipid regulators (**Figure 6b, Supplemental Table S1**). Thus, we concluded that adipocyte Trib1 is regulating plasma lipids through direct regulation by adipose tissue itself.

**Figure 6:**
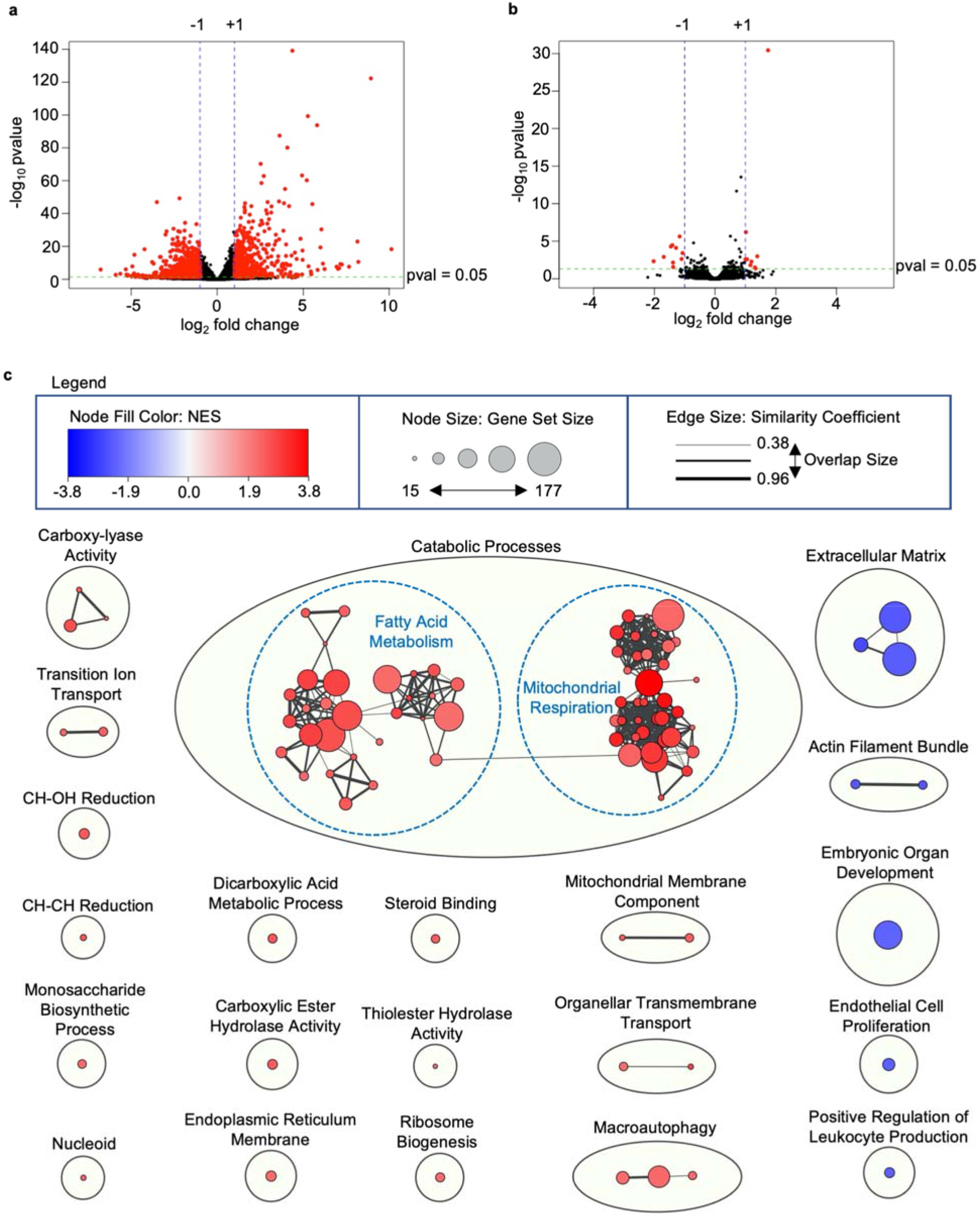
Trib1_ASKO adipocytes have widespread transcriptional changes in mitochondrial and lipid metabolism pathways. **a**, Volcano plot of DESeq2 analysis of RNA-seq data from adipocytes isolated from scWAT from Trib1_fl/fl and Trib1_ASKO mice (*n* = 4). **b**, Volcano plot of DESeq2 analysis of RNA-seq data from hepatocytes from Trib1_fl/fl and Trib1_ASKO mice (*n* = 4). **c**, Cytoscape enrichment plot of Gene Set Enrichment Analysis (GSEA) of differentially expressed adipocyte genes (padj<0.05). Enrichment analysis and clustering were performed as described in the Methods section. Clusters upregulated in Trib1_ASKO samples are shown in red, and clusters upregulated in Trib1_fl/fl samples are shown in blue. Dashed blue outlines indicate larger clusters that were further subclustered manually to facilitate interpretation. NES = normalized enrichment score.

We next ranked differentially expressed genes by signal-to-noise ratio in expression and performed gene set enrichment analysis (GSEA) (**Figure 6c**). Notably, GSEA highlighted a striking enrichment of mitochondrial genes among upregulated genes, including genes coding for proteins in the electron transport chain, mitochondrial ribosomes, and the mitochondrial membrane, suggesting a potential role for mitochondria in the phenotypes we observed. Furthermore, multiple gene sets involved in lipid metabolism were upregulated, pointing towards a role for adipocyte-specific *Trib1* in regulation of lipids through lipid breakdown and metabolism (**Supplemental Table S2**).

### Lipoprotein lipase activity is increased in Trib1_ASKO adipose

Given the importance of adipocytes in TG storage, we next sought to identify the physiological mechanism whereby adipocyte-specific *Trib1* regulates plasma TGs. One mechanism through which adipose contributes to plasma TGs is through the lipolysis of TGs in the lipid droplet and their release as free fatty acids into the bloodstream; subsequently, free fatty acids can be repackaged as TGs and secreted by the liver in the form of VLDL [20]. To assess for an effect on lipolysis by deletion of *Trib1* in adipose, we first measured plasma nonesterified free fatty acids (NEFA) and glycerol, markers of lipolysis, after stimulating lipolysis by fasting mice overnight for 16hr. We found that NEFA and glycerol levels were comparable between Trib1_ASKO and Trib1_fl/fl mice after prolonged fasting (**Supplemental Figure 5a,b**), suggesting that loss of adipocyte *Trib1* does not impact lipolysis rates under stimulation. We also assessed the activation of hormone sensitive lipase (HSL), a key lipolytic driver in adipose that is activated by phosphorylation of key residues, and were unable to detect a difference between phospho-HSL in subcutaneous adipose from Trib1_ASKO and Trib1_fl/fl mice (**Supplemental Figure 5c,d**). Consistent with these observations, VLDL secretion also was unchanged (**Supplemental Figure 5e**), suggesting that the adipose is not providing significantly different loads of fatty acids to the liver.

Given the observed changes in plasma adipokine secretion and the importance of adipose endocrine functions, we next performed an unbiased secretomics experiment to identify differentially secreted proteins from Trib1_ASKO adipose. We incubated scWAT explant tissue from Trib1_fl/fl and Trib1_ASKO mice for 6 hours in serum-free media, and then identified and quantified proteins in the conditioned media via data independent acquisition (DIA) (**Supplemental Table S3**). Consistent with our earlier findings of increased adiponectin and resistin (**Figure 2a,c**), an increase in adiponectin (>50%) and resistin was found in the conditioned media from the ASKO tissue (**Figure 7a**), thus validating our secretomics data. Interestingly, we observed significantly decreased Angptl4 secretion and a trend towards increased Lipoprotein lipase (Lpl) secretion from Trib1_ASKO scWAT explants (**Figure 7a**). Angptl4 is an inhibitor of Lpl, which binds to the endothelium in vasculature and hydrolyzes TGs in circulating lipoproteins to free fatty acids, allowing for their uptake and clearance into tissues, including adipose [21, 22]. In addition to changes in secretion, *Lpl* expression was increased and *Angptl4* expression was decreased significantly in our RNA-seq dataset. The expression of *Lmf1*, which codes for Lipase maturation factor and is important for the proper folding and secretion of Lpl [23], was also notably increased in ASKO adipocytes (**Figure 7b**). Overall, these suggest increased Lpl activity in ASKO adipose tissue. We next measured Lpl activity in adipose tissue extracts from Trib1_fl/fl and Trib1_ASKO mice via cleavage of a fluorescent lipid substrate and found that adipose tissue extracts from both scWAT and VAT from Trib1_ASKO mice demonstrated increased lipase activity (**Figure 7c,d**), likely contributing to increased TG clearance in Trib1_ASKO mice.

**Figure 7:**
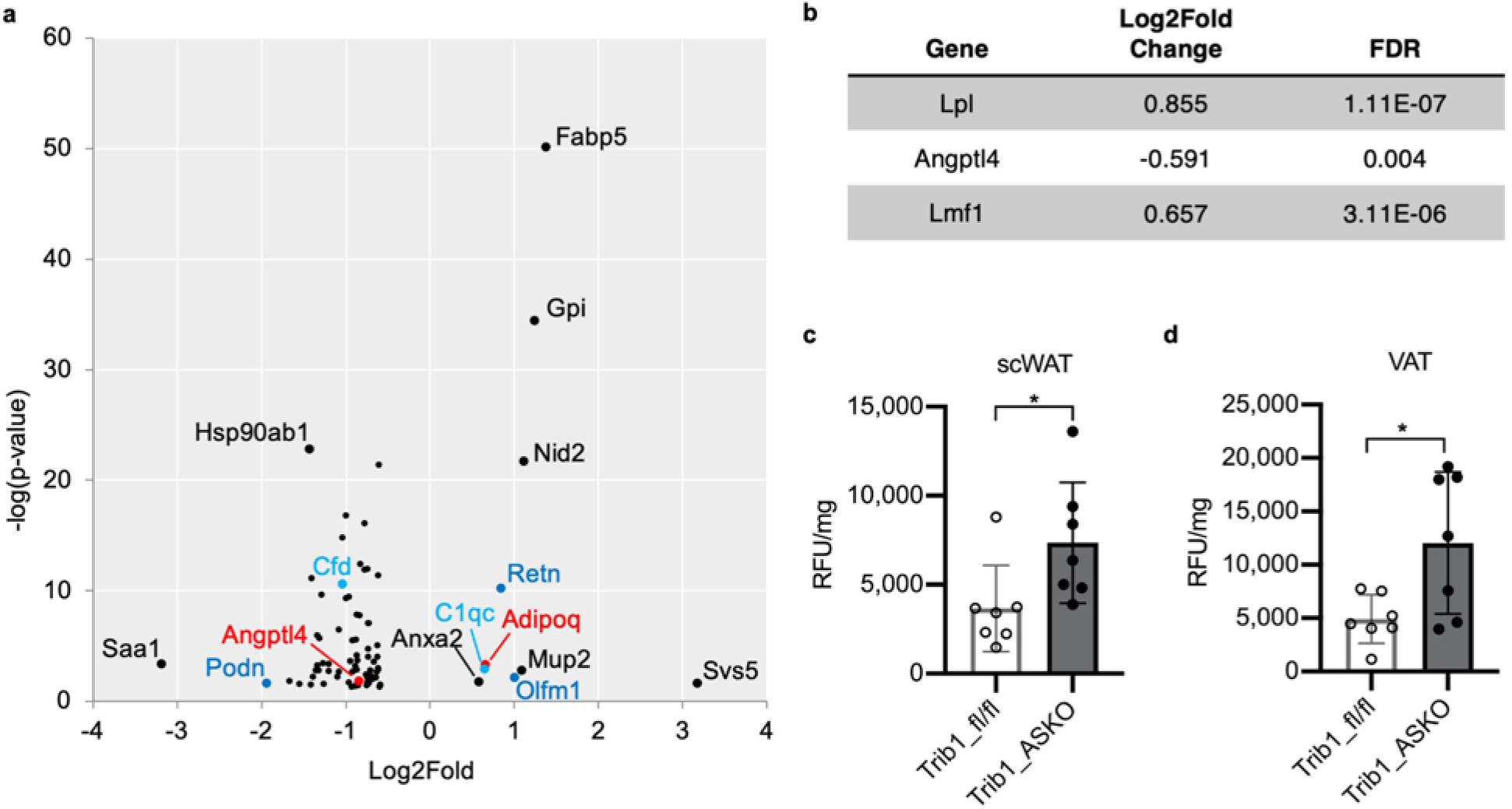
Trib1_ASKO mice have increased adipose tissue lipoprotein lipase activity. **a**, DIA proteomics data of conditioned media from Trib1_ASKO vs. Trib1_fl/fl scWAT explants (*n* = 3). Differential secretion was determined by Spectronaut analysis and results were filtered for secreted proteins (Uniprot keywords). Size and color of datapoints are for facilitating visualization. **b**, DESeq2 results for *Lpl, Angptl4*, and *Lmf1* from RNA-seq of Trib1_fl/fl and Trib1_ASKO adipocytes. **c**,**d**, Lpl activity in scWAT (**c**) and VAT (**d**) extracts (*n* = 5). Data depicted as mean ± s.d. Significance in **c**,**d** determined by Student’ s *t* test (*p <0.05).

## Discussion

Genome-wide association studies have identified SNPs near the *TRIB1* gene that significantly associate with plasma lipids and CAD, and previous work in liver-specific and macrophage-specific mouse models have shown important roles for *Trib1* in plasma lipid regulation as well as in hepatic lipogenesis [7]. An additional GWAS showing an association between the SNPs and circulating adiponectin levels [11] suggested a potential role for adipocyte-specific *TRIB1* in lipid metabolism, and we report here that adipocyte-specific *Trib1* knockout mice have increased plasma adiponectin as well as decreased plasma cholesterol and TGs, thus validating novel roles for adipocyte-specific *Trib1* in both plasma adiponectin and lipid regulation. Interestingly, the reduction in plasma TGs and cholesterol in Trib1_ASKO mice is the opposite of the previously reported liver-specific knock out mice [7], which exhibited increased plasma TC and TG. This suggests tissue-specific roles for *TRIB1* in regulating plasma lipid metabolism, and also highlights the difficulty in determining the causal tissue for associations found in GWAS. Given that there is currently no known functional link between the 8q24 GWAS SNPs and *TRIB1* expression or function [24], further functional genomic studies will be required to understand if and how these SNPs contribute to tissue-specific TRIB1 function.

TRIB1 is one of three mammalian homologs of the Tribbles pseudokinase that was first discovered in *Drosophila* [17]. These proteins bear homology to serine/threonine kinases, but lack key catalytic residues that render them unable to catalyze phosphorylation. Instead, they are best understood to function as scaffolding proteins that bring other proteins into proximity with each other to mediate signaling events [17]. One of the best understood molecular functions for TRIB1 is its role in mediating the ubiquitination and degradation of the transcription factor C/EBPα by bringing it into proximity of the COP1 E3 ubiquitin ligase. Tribbles-mediated regulation of C/EBPα protein levels has been shown to be an important function in several models, including as a causal mechanism for the hepatic lipogenesis phenotype in LSKO mice [7], for myeloid cell proliferation in the context of leukemia [18], and in oogenesis in drosophila [25]. We report here that Trib1_ASKO adipocytes also exhibit increased C/EBPα protein levels in the absence of any change in gene expression, and that C/EBPα protein levels are normalized between control and Trib1_ASKO SVF-derived adipocytes under conditions of proteasomal inhibition. Thus, we provide evidence that adipocyte Trib1 also regulates C/EBPα through proteasomal degradation.

C/EBPα is a critical regulator of adipocyte differentiation [26]. However, despite increased C/EBPα protein, we interestingly did not observe any differences in adiposity or adipose morphology in Trib1_ASKO mice, or in the *in vitro* differentiation of adipose stem cells from the ASKO mice. This could be a result of the *Adipoq* promoter-driven Cre, which is induced late in the process of adipocyte differentiation, thus making our mouse model a post-differentiation knockout of adipocyte Trib1. A previous report showed that Trib1 overexpression can inhibit differentiation of 3T3-L1 cells [27], providing precedent for a role for Trib1 in adipogenesis. Further studies utilizing a different Cre transgene would be required to determine if Trib1 has a similar role in regulating adipogenesis *in vivo*.

We also found that Trib1_ASKO adipocytes have both increased cellular levels of adiponectin and increased secretion of adiponectin, with the former likely driving the latter. We observed no change in *Adipoq* gene expression via repeated qPCR measurements in multiple *ex vivo* cell culture experiments, whole adipose tissue, and isolated adipocytes from Trib1_ASKO mice. Thus, the increase in adiponectin protein comes in the absence of any reliable change in *Adipoq* gene expression, suggesting a post-transcriptional mechanism of regulation. We will note, however, that *Adipoq* expression was surprisingly increased in Trib1_ASKO mice in our RNA-seq dataset (padj = 0.011, fold change = 1.33). This raises the possibility that increased protein levels of C/EBPα, which is a well-known transcriptional regulator of adiponectin expression [19], or a different unknown transcription factor is driving a small increase in *Adipoq* gene expression that qPCR is not sensitive enough to reliably measure. We note though that while MG132 treatment of SVF-derived adipocytes increases C/EBPα protein levels, it actually decreases the secretion of adiponectin in wild-type SVF-derived adipocytes (**Figure 5d)**. Thus, while Trib1 certainly appears to regulate C/EBPα in adipose, we propose this is a separate mechanism from the one governing Trib1 regulation of adiponectin. The exact nature of the relationship between Trib1 and cellular adiponectin levels remains to be determined.

As noted, Trib1_ASKO mice exhibit decreased plasma TC and TG levels. We subsequently determined that these mice also have increased adipose Lpl activity, likely driving increased uptake of plasma TGs into adipocytes and contributing to the reduction in plasma TG. This might be expected to drive an increase in adipocyte size, which we did not observe in Trib1_ASKO mice. However, GSEA of our RNA-seq data revealed upregulation of genes encoding mitochondrial components, suggestive of increased mitochondrial activity. A resulting increase in energy expenditure could potentially explain the lack of an adipocyte size phenotype in Trib1_ASKO mice despite increased adipose tissue Lpl activity and presumed fatty acid uptake. Notably, C/EBPα regulates genes involved in lipid metabolism in adipose tissue and is a known transcriptional regulator of *Lpl* [28, 29], and polymorphisms in C/EBPα have also been found to associate with plasma TG levels in humans [29]. It is thus possible that increased C/EBPα protein levels in Trib1_ASKO adipose may contribute to this lipid phenotype. There is some precedent that the increased Lpl activity in the Trib1_ASKO could contribute to the observed decrease in plasma cholesterol. Multiple studies using transgenic Lpl animal models [30-32] as well as Angptl4 knockout or transgenic mice [33, 34] demonstrate that increased Lpl activity protects from diet-induced hypercholesterolemia and decreases plasma LDL-C levels, though the effects are not as robust as effects on plasma TGs. Mechanistically, lipolysis-mediated reductions in TGs in VLDL particles have been proposed to facilitate enhanced clearance of the resulting remnant particles via receptors such as the LDLR [35, 36]. Further studies are necessary to determine if increased LPL activity is responsible for the cholesterol phenotype in Trib1_ASKO mice.

Adiponectin has many well-studied roles in regulating cardiometabolic traits, including lipid metabolism and coronary artery disease [13, 15]. Thus, an important outstanding question is whether adiponectin is driving the lipid phenotypes observed in the Trib1_ASKO mice. Adiponectin’ s role in insulin sensitization is perhaps its most widely recognized physiological effect [37, 38], and indeed we found that Trib1_ASKO mice demonstrated improved glucose tolerance compared to Trib1_fl/fl mice when placed on high-fat diet. However, a role for adiponectin in regulating plasma cholesterol is less clear. One previous report using adiponectin transgenic mice with 10-fold increased adiponectin levels found decreased cholesterol in those mice [39]. In humans, numerous epidemiological studies have been conducted looking at associations between adiponectin and plasma LDL-cholesterol, yet many of these are conflicting or inconclusive [13]. Epidemiological studies and genetic mouse models do provide clear support for a role for adiponectin in TG and VLDL metabolism. In particular, plasma adiponectin correlates with decreased plasma TGs and increased HDL-C in humans [40], and adiponectin transgenic mice with 3-fold increased plasma adiponectin levels have increased Lpl expression and activity in adipose tissue as well as increased TG clearance [41]. Thus, although the adiponectin phenotype in Trib1_ASKO mice is mild (∼20-30% increase) compared to transgenic mouse models, it is possible that the increased adipose tissue Lpl activity we observe in Trib1_ASKO mice is secondary to increased adiponectin levels. Further studies will be necessary to determine if the adiponectin phenotype is required for the observed changes in Trib1_ASKO plasma lipids.

Overall, our studies show that adipocyte-specific *Trib1* is a negative regulator of adiponectin secretion, and that this appears to be through a C/EBPα-independent mechanism. Furthermore, we show that adipocyte-specific *Trib1* regulates plasma lipids in a direction opposite to that of the previously studied LSKO model, and that regulation of TG clearance via adipose Lpl in part explains the decreased plasma TG levels in ASKO mice. In contrast to hepatic Trib1, these data suggest a therapeutically beneficial effect of reduced adipocyte *Trib1* activity, underscoring the continued importance of further studies on *Trib1* and the 8q24 lipid and CAD GWAS locus.

## Methods

### Animals

The previously reported Trib1_fl/fl mice (Bauer et al, JCI 2015) were bred in house. *Adipoq*-Cre mice (stock#010803) and Ldlr KO mice (stock#002207) were obtained from Jackson Labs. Mice were fed ad-libitum on chow diet unless otherwise noted. All experiments were performed when mice were 8-12 weeks old. Mice were fasted for 4 hr prior to collecting plasma samples, unless stated otherwise. Blood was collected retro-orbitally and spun at 10,000 rpm for 7 min. Fasting cholesterol and TGs were measured via plate assay using Infinity reagents (Fisher TR13421 and TR22421), and adipokine levels were measured via ELISA (adiponectin: Millipore EZ-MADPK, leptin: Millipore EZML-82K, resistin: R&D MRSN00). For HFD experiments, mice were placed on 45% kcal HFD (Research Diets D12451) starting at 8-12 weeks of age. Plasma was collected retro-orbitally at 0, 4, 8, and 12 weeks of HFD, and glucose tolerance testing performed at 0, 6, and 12 weeks of HFD as previously described [42]. For fasting/refeeding experiments, mice were fasted overnight for 16 hr and then fed ad-libitum with chow diet for 3hr. To measure *in vivo* TG secretion, plasma triglycerides were measured in 4 hr-fasted mice 30, 60, 120, and 180 min after i.p. injection of 1mg pluronic (P407) per gram mouse body weight. All *in vivo* studies described here were approved by Columbia University’ s Institutional Animal Care and Use Committee prior to commencement.

### FPLC analysis of pooled plasma

200 μl of pooled plasma from gender and genotype matched mice (n = 4-9) was loaded onto a Superose 6 column (GE Healthcare) calibrated with elution buffer (0.15 M NaCl, 1 mM EDTA). The lipoproteins were eluted in a total of 20 mL elution buffer in 0.5 mL fractions at a rate of 0.3 mL/min. The cholesterol and TG content of each fraction was determined by plate assay.

### Western Blot Analysis

Tissues or cells were lysed and homogenized in RIPA buffer supplemented with 1x Halt Protease and Phosphatase inhibitor (Fisher Scientific PI78444). The lysate was centrifuged at 12,000 xg for 15 min at 4 oC to clarify the protein prep from cellular debris and lipids. ∼30 ug protein was loaded onto 10% bis-tris SDS-PAGE gel and transferred onto a nitrocellulose membrane. The membrane was blocked in either 5% milk or BSA (for phospho-protein analysis) and incubated in the appropriate primary antibody (adiponectin (R&D AF1119), Trib1 (Millipore 09-126), Flag (Sigma F7425), Tubulin (CST 3873S), C/EBPα (CST 2295S), Beta-actin (Santa Cruz sc-81178), Hsl, pHsl565, pHsl563, and pHsl660 (CST 8334T)) overnight. Protein was detected using a secondary HRP-linked antibody and Luminata Classico Western HRP Substrate (Millipore WBLUC0020). To reprobe membranes, membranes were incubated in stripping buffer (Fisher Scientific PI21059) for 15 min before reblocking.

### qPCR analysis

RNA from tissues and cells were isolated using the RNeasy Mini kit (Qiagen). cDNA was synthesized using the High-Capacity cDNA Reverse Transcription Kit (Applied Biosystems). qPCR was performed using predesigned Taqman probes from Thermo Fisher Scientific. Gene expression data was normalized to *Gapdh* and presented as fold change relative to the Trib1_fl/fl control group (exceptions indicated in the figure legend).

### Microscopy

For adipose tissue histology, scWAT samples (<4 mm thick) from Trib1_fl/fl and Trib1_ASKO mice were fixed in 4% PFA for 24 hr. The tissues were then embedded in paraffin, sectioned at 7 μm, and H&E stained. For each mouse, 4 sections at 70 μm intervals were imaged on a Nikon Eclipse Ti microscope with the 40x objective and analyzed using the Adiposoft ImageJ plugin (parameters: minimum diameter = 10 μm, maximum diameter = 100 μm). For Oil Red O staining, cells were fixed in 4% PFA for 15 min and then placed in 0.3% w/v Oil Red O in 60% isopropanol for 30 min. The cells were washed 5X in distilled H2O, and then imaged with the 20x objective.

### RNA-seq of adipocytes and hepatocytes

8–12-week-old male mice were euthanized and perfused with PBS after a 4 hr fast. Subcutaneous inguinal fat pads from individual mice were harvested, minced, and then placed in 6 mL digestion media (0.14 U/mL Liberase TM, 50 U/mL DNAse I, 20 mg/mL BSA in DMEM) for 1 hr at 37 oC, shaking at 250 rpm. The tissue prep was then filtered through a 100 μm cell strainer, and spun at 300 xg for 10 min. The floating white layer was collected as the adipocyte fraction and placed in 1mL Qiazol. RNA was then isolated using the RNeasy Lipid Tissue Mini Kit (Qiagen). Livers from the same mice were harvested and homogenized in Trizol, and RNA was isolated via chloroform extraction. RNA quality was assessed via BioAnalyzer before being submitted to the core for bulk, paired-end RNA-sequencing (NextSeq 500). Reads were aligned using STAR and featurecounts, and differential expression analysis was performed using the DESeq2 package. Differentially expressed genes (padj < 0.050) were ranked by Signal-to-noise ratio of median normalized counts and analyzed by GSEA using the Gene Ontology gene sets (c5.go.v7.2.symbols.gmt) from MSigDB, using gene set size ≤ 200 and 1000 permutations of the gene sets to determine enrichment score. Cytoscape enrichment plots were constructed from GSEA results using FDR < 0.01, and a combined coefficient > 0.375 with combined constant 0.5 as described in [43]. Nodes were clustered using the MCL clustering algorithm in the Autoannotate Cytoscape App. Annotations of clusters were manually curated.

### SVF Generation and Differentiation

Subcutaneous inguinal fat pads from 3-5 mice of the same gender and genotype were combined and minced in digestion buffer (L-15 Leibovitz media, 1.5% BSA, 1% Pen/Strep, 10 U/mL DNaseI, 480 U/mL Hyaluronidase, 0.14 U/mL Liberase TM). Tissue was allowed to dissociate in digestion buffer for 1 hr at 37 oC, shaking at 250 rpm. The tissue prep was then filtered through a 100 μM cell strainer and spun at 300 xg, 4oC, for 10 min. The pellet was saved and resuspended in 10 mL culture medium (DMEM, 10% FBS, 1% Pen/Strep, 2 mM L-Glut). The cells were spun at 300 xg, 4 oC, for 10 min, and resuspended in 5 mL culture medium supplemented with 1 μg/mL insulin before seeding. Media was changed every 2 – 3 days until the cells were >95% confluent. Differentiation was initiated with a cocktail including 10% FBS, 1% Pen/Strep, 5 μg/mL insulin, 1 μM Rosiglitazone, 1 μM Dexamethasone, and 250 μM IBMX in DMEM/F12. After 48 hr, cells were maintained in DMEM/F12 supplemented with only 10% FBS, 1% Pen/Strep, 5 μg/mL insulin, and 1 μM Rosiglitazone. Experiments were started after day 7 of differentiation.

### Global quantitative proteomics of Explant secretomics

Mice were euthanized and perfused with PBS before dissection of subcutaneous adipose fat pads. 50 mg of tissue was placed into 1mL of warm, serum-free DMEM in a 12 well plate and pinned down with transwell insert. The media was collected after 6hr and protein was precipitated using methanol. DIA (Data independent acquisition) based proteomics was used. In brief, protein precipitated pellets were resuspended in SDC lysis buffer [44] (1% SDC, 10 mM TCEP, 40 mM CAA and 100 mM Tris-HCl pH 8.5) and boiled for 10 min at 95°C, 1500 rpm to denature and reduce and alkylate cysteins, followed by sonication in a water bath, cooled down to room temperature. Protein concentration was estimated by BCA measurement and 20 µg were further processed for overnight digestion by adding LysC and trypsin in a 1:50 ratio (µg of enzyme to µg of protein) at 37° C and 1500 rpm. Peptides were acidified by adding 1% TFA, vortexed, and subjected to StageTip clean-up via SDB-RPS. 20 µg of peptides were loaded on two 14-gauge StageTip plugs. Peptides were washed two times with 200 µL 1% TFA 99% ethyl acetate followed 200 µL 0.2% TFA/5%ACN in centrifuge at 3000 rpm, followed by elution with 60 µL of 1% Ammonia, 50% ACN into eppendorf tubes and dried at 60°C in a SpeedVac centrifuge. Peptides were resuspended in 10 µL of 3% acetonitrile/0.1% formic acid and injected on Thermo Scientific™ Orbitrap Fusion™ Tribrid™ mass spectrometer with DIA method [45] for peptide MS/MS analysis. The UltiMate 3000 UHPLC system (Thermo Scientific) and EASY-Spray PepMap RSLC C18 50 cm x 75 μm ID column (Thermo Fisher Scientific) coupled with Orbitrap Fusion (Thermo) were used to separate fractioned peptides with a 5-30% acetonitrile gradient in 0.1% formic acid over 127 min at a flow rate of 250 nL/min. After each gradient, the column was washed with 90% buffer B for 5 min and re-equilibrated with 98% buffer A (0.1% formic acid, 100% HPLC-grade water) for 40min. Survey scans of peptide precursors were performed from 350-1200 *m/z* at 120K FWHM resolution (at 200 *m/z*) with a 1 × 10^6^ ion count target and a maximum injection time of 60 ms. The instrument was set to run in top speed mode with 3 s cycles for the survey and the MS/MS scans. After a survey scan, 26 m/z DIA segments will be acquired at from 200-2000 *m/z* at 60K FWHM resolution (at 200 *m/z*) with a 1 × 10^6^ ion count target and a maximum injection time of 118 ms. HCD fragmentation was applied with 27% collision energy and resulting fragments were detected using the rapid scan rate in the Orbitrap. The spectra were recorded in profile mode. DIA data were analyzed with directDIA 2.0 (Deep learning augmented spectrum-centric DIA analysis) in Spectronaut Pulsar X, a mass spectrometer vendor independent software from Biognosys. The default settings were used for targeted analysis of DIA data in Spectronaut except the decoy generation was set to “mutated”. The false discovery rate (FDR) will be estimated with the mProphet approach and set to 1% at peptide precursor level and at 1% at protein level.

Results obtained from Spectronaut were further analyzed using the Spectronaut statistical package. Significantly changed protein abundance was determined by un-paired t-test with a threshold for significance of p < 0.05 (permutation-based FDR correction) and 0.58 log2FC.

### Cloning and Lentivirus Production

Lentiviral constructs for tetracycline-inducible expression of proteins (mTrib1 and eGFP) in adipocytes for overexpression experiments were cloned by first introducing a 3xFlagHA tag at the C-terminal end of each protein. The fusion proteins were then cloned into the pEN-TTMCS entry vector to introduce a tight TRE promoter and subsequently cloned into the pSLIK-neo lentiviral plasmid via Gateway cloning. mTrib1 and eGFP were also cloned into the pLentiCMVPuroDEST lentiviral vector for constitutive overexpression under the CMV promoter. To produce the virus, 5 × 10^6^ 293T cells were seeded in T75 flasks and transfected with 2 μg MD2G, 3 μg Pax2, and 5 μg lentiviral construct with 30 μL Fugene 6 (Promega) the following day. The media was changed the day after transfection, and the viral supernatant was collected and pooled after 24hr and 48hr. The supernatant was filtered through a 0.45 μm filter, aliquoted, and stored at −80oC until use.

### Adipocyte Cell culture

3T3-L1 cells were purchased from ATCC and cultured in DMEM supplemented with 10% FBS, 1 mM Sodium Pyruvate, and 1% Pen/Strep. Cells were tested for mycoplasma every three months. To differentiate 3T3-L1 cells to adipocytes, cells were induced with growth media supplemented with 1 μM Dexamethasone, 0.5 mM IBMX, and 1 μg/mL Insulin for 48 hr, and then maintained in growth media supplemented with only 1 μg/mL insulin. Experiments were typically performed on cells 7 – 9 days after differentiation induction. Stable doxycycline-inducible 3T3-L1 cells were generated by transducing cells with lentivirus at an MOI ∼100, followed by selection with 1.5 μg/mL puromycin. Conditioned media was collected in OptiMEM I reduced serum media.

### Fluorescent LPL Assay

The Lpl activity assay was adapted from Basu et. al [46]. Briefly, adipose tissue was minced in 5 μl x mg tissue weight volume in tissue incubation buffer (PBS, 2 mg/ml FA-free BSA, 5 U/mL heparin), incubated for 1 hr in a 37 oC shaker, and centrifuged at 3,000 rpm for 15 min at 4 oC. The clarified supernatant was placed in fresh tubes and diluted 1:10 in tissue incubation buffer. 4 μl of lysate was placed in duplicate in a black-walled 96-well plate, and 100 μL reaction buffer (0.15 M NaCl, 20 mM Tris-HCl pH 8.0, 0.0125% Zwittergent, 1.5% FA-free BSA, 0.62 μM EnzChek (Invitrogen E33955)) was added to each well. The reaction was allowed to incubate 20 min at 37 oC, and was then read at an excitation of 485 and emission of 515. A blank RFU value was subtracted from all experimental RFU values, and the resulting values were reported.

### Statistics

GraphPad Prism 8 was used to graph data and to perform parametric 2-tailed Student’ s t tests and 1-and 2-way ANOVA analyses with multiple correction using either Dunnett’ s, Sidak’ s, or Tukey’ s method as indicated in the figure legends.

## Acknowledgements

These studies were funded by R01HL141745 (R.C.B) from the NIH/NHLBI and Scientific Development Grant 16SDG31180039 (R.C.B) from the American Heart Association. Additionally, E.E.H was supported by a Ruth L. Kirschstein Individual Predoctoral F30 NRSA (F30HL146076-01A1) from the NIH/NHLBI.

## Author Contributions

R.C.B conceived the project, designed the experiments, supervised analyses and edited the manuscript. E.E.H. performed the majority of the experiments and data analysis, and wrote the first draft of the manuscript and edited subsequent versions. G.I.Q helped establish the mouse colony, assisted with SVF isolation, and performed related molecular biology (i.e. cloning, western blots). R.L. performed western blotting for adipose lipolysis proteins and ELISA analysis. C.X. performed the DESeq2 analysis of the RNA-seq data. A.H. performed initial FPLC analysis and assisted with all FPLC analysis. R.I. performed SVF isolation and cloning of viral vectors. J.C. managed the animal colony and assisted with insulin trait experiments. R.K.S. performed and analyzed the secretomics MS experiment. All the authors read and approved the manuscript.

## Competing Interests Statement

The Authors declare no competing interests.

## Data Availability Statement

GEO accession numbers for RNA-seq data will be available prior to publication. Full list of identified proteins and differentially secreted proteins from DIA secretomics (Figure 7a) is available in supplemental table S3. Other data that support the findings of this study are available from the corresponding author upon reasonable request.

## Notes

### Competing Interest Statement

The authors have declared no competing interest.

## References

1. Willer, C.J., et al., Newly identified loci that influence lipid concentrations and risk of coronary artery disease. Nat Genet, 2008. 40(2): p. 161–9.

2. Kathiresan, S., et al., Six new loci associated with blood low-density lipoprotein cholesterol, high-density lipoprotein cholesterol or triglycerides in humans. Nat Genet, 2008. 40(2): p. 189–97.

3. Teslovich, T.M., et al., Biological, clinical and population relevance of 95 loci for blood lipids. Nature, 2010. 466(7307): p. 707–13.

4. Consortium, I.K.C., Large-scale gene-centric analysis identifies novel variants for coronary artery disease. PLoS Genet, 2011. 7(9): p. e1002260.

5. Consortium, C.A.D., et al., Large-scale association analysis identifies new risk loci for coronary artery disease. Nat Genet, 2013. 45(1): p. 25–33.

6. Willer, C.J., et al., Discovery and refinement of loci associated with lipid levels. Nat Genet, 2013. 45(11): p. 1274–1283.

7. Bauer, R.C., et al., Tribbles-1 regulates hepatic lipogenesis through posttranscriptional regulation of C/EBPalpha. J Clin Invest, 2015. 125(10): p. 3809–18.

8. Burkhardt, R., et al., Trib1 is a lipid- and myocardial infarction-associated gene that regulates hepatic lipogenesis and VLDL production in mice. J Clin Invest, 2010. 120(12): p. 4410–4.

9. Johnston, J.M., et al., Myeloid Tribbles 1 induces early atherosclerosis via enhanced foam cell expansion. Sci Adv, 2019. 5(10): p. eaax9183.

10. Chambers, J.C., et al., Genome-wide association study identifies loci influencing concentrations of liver enzymes in plasma. Nat Genet, 2011. 43(11): p. 1131–8.

11. Dastani, Z., et al., Novel loci for adiponectin levels and their influence on type 2 diabetes and metabolic traits: a multi-ethnic meta-analysis of 45,891 individuals. PLoS Genet, 2012. 8(3): p. e1002607.

12. Lihn, A.S., S.B. Pedersen, and B. Richelsen, Adiponectin: action, regulation and association to insulin sensitivity. Obes Rev, 2005. 6(1): p. 13–21.

13. Izadi, V., E. Farabad, and L. Azadbakht, Epidemiologic evidence on serum adiponectin level and lipid profile. Int J Prev Med, 2013. 4(2): p. 133–40.

14. Xu, A., et al., The fat-derived hormone adiponectin alleviates alcoholic and nonalcoholic fatty liver diseases in mice. J Clin Invest, 2003. 112(1): p. 91–100.

15. Pischon, T., et al., Plasma adiponectin levels and risk of myocardial infarction in men. JAMA, 2004. 291(14): p. 1730–7.

16. Ostertag, A., et al., Control of adipose tissue inflammation through TRB1. Diabetes, 2010. 59(8): p. 1991–2000.

17. Eyers, P.A., K. Keeshan, and N. Kannan, Tribbles in the 21st Century: The Evolving Roles of Tribbles Pseudokinases in Biology and Disease. Trends Cell Biol, 2017. 27(4): p. 284–298.

18. Dedhia, P.H., et al., Differential ability of Tribbles family members to promote degradation of C/EBPalpha and induce acute myelogenous leukemia. Blood, 2010. 116(8): p. 1321–8.

19. Christy, R.J., et al., Differentiation-induced gene expression in 3T3-L1 preadipocytes: CCAAT/enhancer binding protein interacts with and activates the promoters of two adipocyte-specific genes. Genes Dev, 1989. 3(9): p. 1323–35.

20. Duncan, R.E., et al., Regulation of lipolysis in adipocytes. Annu Rev Nutr, 2007. 27: p. 79–101.

21. Dijk, W. and S. Kersten, Regulation of lipoprotein lipase by Angptl4. Trends Endocrinol Metab, 2014. 25(3): p. 146–55.

22. Goldberg, I.J. and M. Merkel, Lipoprotein lipase: physiology, biochemistry, and molecular biology. Front Biosci, 2001. 6: p. D388–405.

23. Doolittle, M.H., N. Ehrhardt, and M. Peterfy, Lipase maturation factor 1: structure and role in lipase folding and assembly. Curr Opin Lipidol, 2010. 21(3): p. 198–203.

24. Jadhav, K.S. and R.C. Bauer, Trouble With Tribbles-1. Arterioscler Thromb Vasc Biol, 2019. 39(6): p. 998–1005.

25. Rorth, P., K. Szabo, and G. Texido, The level of C/EBP protein is critical for cell migration during Drosophila oogenesis and is tightly controlled by regulated degradation. Mol Cell, 2000. 6(1): p. 23–30.

26. Farmer, S.R., Transcriptional control of adipocyte formation. Cell Metab, 2006. 4(4): p. 263–73.

27. Naiki, T., et al., TRB2, a mouse Tribbles ortholog, suppresses adipocyte differentiation by inhibiting AKT and C/EBPbeta. J Biol Chem, 2007. 282(33): p. 24075–82.

28. Madsen, M.S., et al., Peroxisome proliferator-activated receptor gamma and C/EBPalpha synergistically activate key metabolic adipocyte genes by assisted loading. Mol Cell Biol, 2014. 34(6): p. 939–54.

29. Olofsson, L.E., et al., CCAAT/enhancer binding protein alpha (C/EBPalpha) in adipose tissue regulates genes in lipid and glucose metabolism and a genetic variation in C/EBPalpha is associated with serum levels of triglycerides. J Clin Endocrinol Metab, 2008. 93(12): p. 4880–6.

30. Shimada, M., et al., Suppression of diet-induced atherosclerosis in low density lipoprotein receptor knockout mice overexpressing lipoprotein lipase. Proc Natl Acad Sci U S A, 1996. 93(14): p. 7242–6.

31. Fan, J., et al., Overexpression of lipoprotein lipase in transgenic rabbits inhibits diet-induced hypercholesterolemia and atherosclerosis. J Biol Chem, 2001. 276(43): p. 40071–9.

32. Walton, R.G., et al., Increasing adipocyte lipoprotein lipase improves glucose metabolism in high fat diet-induced obesity. J Biol Chem, 2015. 290(18): p. 11547–56.

33. Koster, A., et al., Transgenic angiopoietin-like (angptl)4 overexpression and targeted disruption of angptl4 and angptl3: regulation of triglyceride metabolism. Endocrinology, 2005. 146(11): p. 4943–50.

34. Aryal, B., et al., Absence of ANGPTL4 in adipose tissue improves glucose tolerance and attenuates atherogenesis. JCI Insight, 2018. 3(6).

35. Aviram, M., E.L. Bierman, and A. Chait, Modification of low density lipoprotein by lipoprotein lipase or hepatic lipase induces enhanced uptake and cholesterol accumulation in cells. J Biol Chem, 1988. 263(30): p. 15416–22.

36. Sehayek, E., U. Lewin-Velvert, T. Chajek-Shaul, and S. Eisenberg, Lipolysis exposes unreactive endogenous apolipoprotein E-3 in human and rat plasma very low density lipoprotein. J Clin Invest, 1991. 88(2): p. 553–60.

37. Tilg, H. and A.R. Moschen, Adipocytokines: mediators linking adipose tissue, inflammation and immunity. Nat Rev Immunol, 2006. 6(10): p. 772–83.

38. Yanai, H. and H. Yoshida, Beneficial Effects of Adiponectin on Glucose and Lipid Metabolism and Atherosclerotic Progression: Mechanisms and Perspectives. Int J Mol Sci, 2019. 20(5).

39. Bauche, I.B., et al., Overexpression of adiponectin targeted to adipose tissue in transgenic mice: impaired adipocyte differentiation. Endocrinology, 2007. 148(4): p. 1539–49.

40. Christou, G.A. and D.N. Kiortsis, Adiponectin and lipoprotein metabolism. Obes Rev, 2013. 14(12): p. 939–49.

41. Combs, T.P., et al., A transgenic mouse with a deletion in the collagenous domain of adiponectin displays elevated circulating adiponectin and improved insulin sensitivity. Endocrinology, 2004. 145(1): p. 367–83.

42. Lagor, W.R., et al., Deletion of murine Arv1 results in a lean phenotype with increased energy expenditure. Nutr Diabetes, 2015. 5: p. e181.

43. Reimand, J., et al., Pathway enrichment analysis and visualization of omics data using g:Profiler, GSEA, Cytoscape and EnrichmentMap. Nat Protoc, 2019. 14(2): p. 482–517.

44. Kulak, N.A., et al., Minimal, encapsulated proteomic-sample processing applied to copy-number estimation in eukaryotic cells. Nat Methods, 2014. 11(3): p. 319–24.

45. Bruderer, R., et al., Extending the limits of quantitative proteome profiling with data-independent acquisition and application to acetaminophen-treated three-dimensional liver microtissues. Mol Cell Proteomics, 2015. 14(5): p. 1400–10.

46. Basu, D., J. Manjur, and W. Jin, Determination of lipoprotein lipase activity using a novel fluorescent lipase assay. J Lipid Res, 2011. 52(4): p. 826–32.

