## Supplemental Figures for "Adipocyte Tribbles1 Regulates Plasma Adiponectin and Plasma Lipids in Mice"

**Figure S1:**

**a**

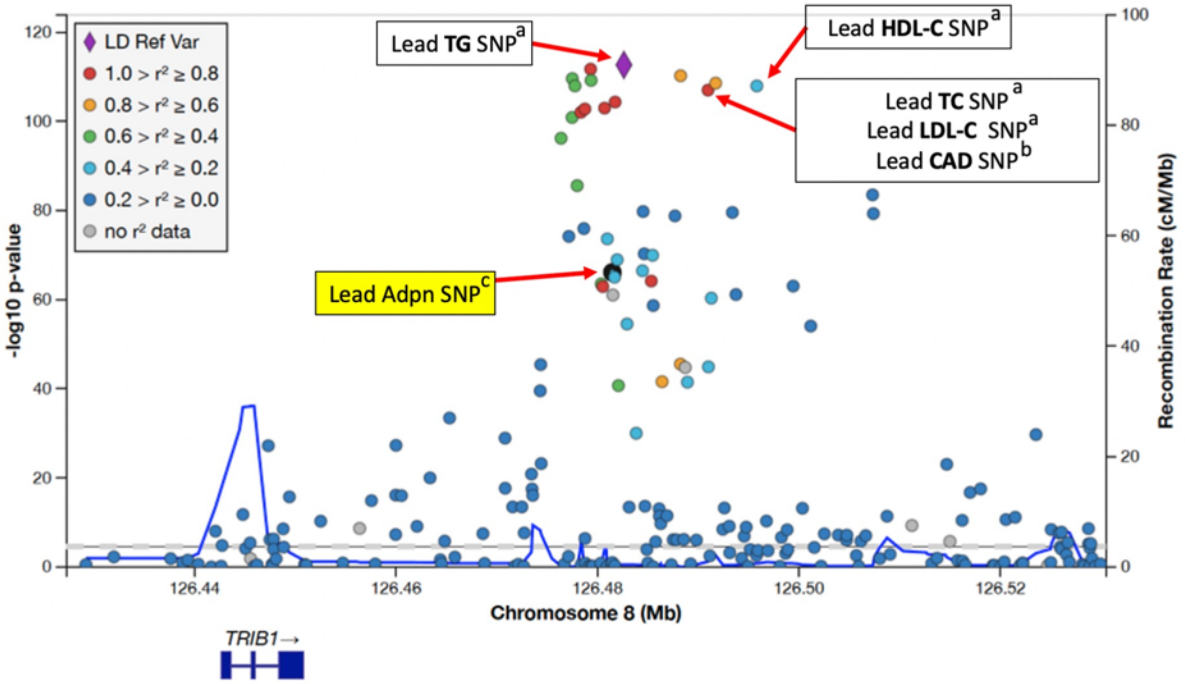


**b**

| SNP | Genomic Location | TC p-value | TG p-value | Adpn p-value |
|---|---|---|---|---|
| rs2980879 | chr8:126481475 | 2.42e-25 | 8.019e-67 | 1.1e-08 |
| rs2954022 | chr8:126482621 | 4.354e-61 | 2.23e-113 | 7.51e-06 |
| rs2954029 | chr8:126490972 | 2.421e-65 | 1.02e-107 | 1.76e-05 |

### Figure S2

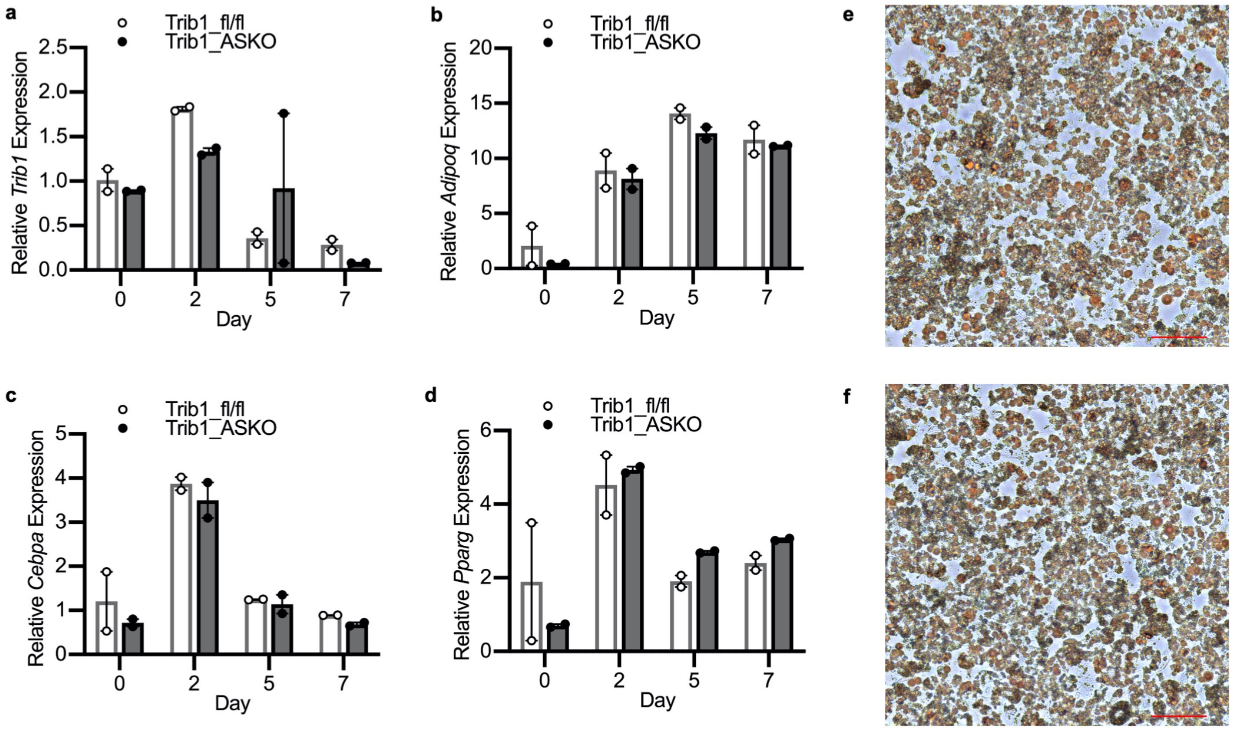
a
Trib1_fl/fl
Trib1_ASKO
Relative Trib1 Expression
2.5
2.0
1.5
1.0
0.5
0.0
0
2
5
7
Day
b
Trib1_fl/fl
Trib1_ASKO
Relative Adipoq Expression
20
15
10
5
0
0
2
5
7
Day
c
Trib1_fl/fl
Trib1_ASKO
Relative Cebpa Expression
5
4
3
2
1
0
0
2
5
7
Day
d
Trib1_fl/fl
Trib1_ASKO
Relative Pparg Expression
6
4
2
0
0
2
5
7
Day
e
f

**Figure S3**

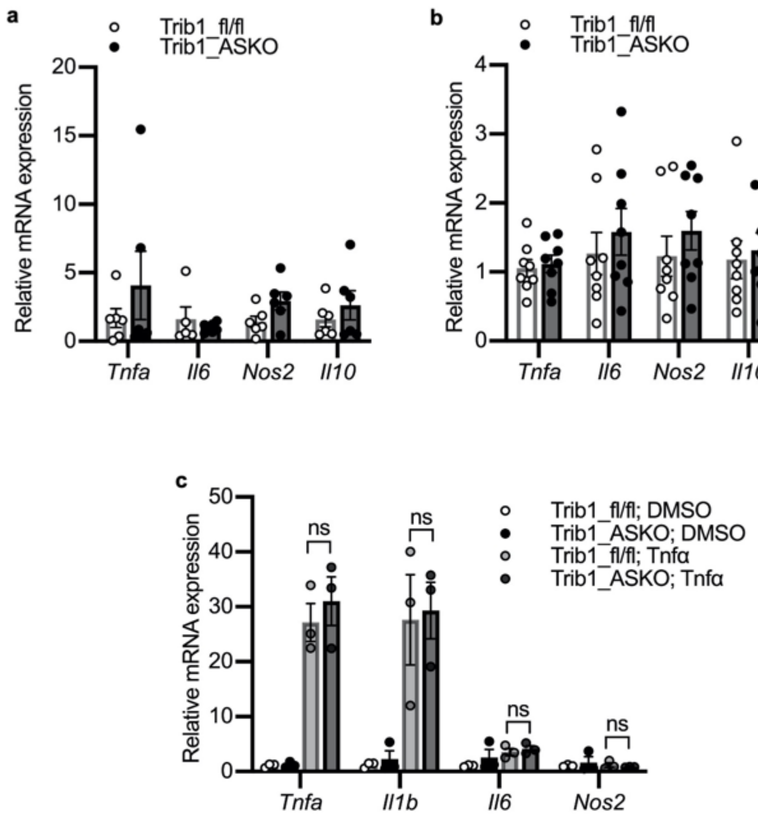
a
Trib1_fl/fl
Trib1_ASKO
Relative mRNA expression
20
15
10
5
0
Tnfa
Il6
Nos2
Il10
b
Trib1_fl/fl
Trib1_ASKO
Relative mRNA expression
4
3
2
1
0
Tnfa
Il6
Nos2
Il10
c
Relative mRNA expression
50
40
30
20
10
0
ns
ns
ns
ns
Trib1_fl/fl; DMSO
Trib1_ASKO; DMSO
Trib1_fl/fl; Tnfα
Trib1_ASKO; Tnfα
Tnfa
Il1b
Il6
Nos2

**Figure S4:**

a

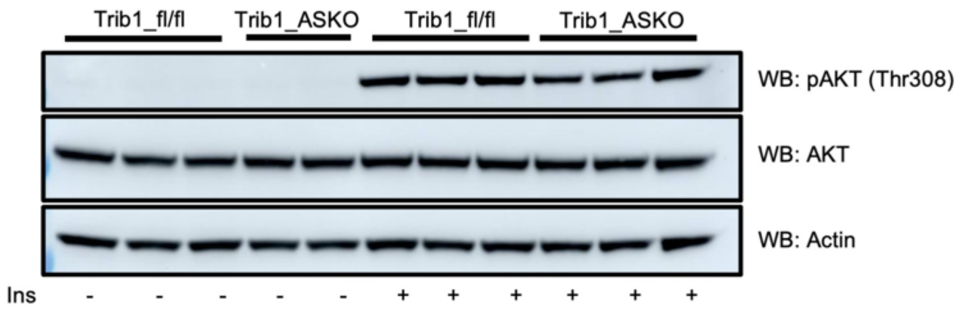

Trib1_fl/fl
Trib1_ASKO
Trib1_fl/fl
Trib1_ASKO
WB: pAKT (Thr308)
WB: AKT
WB: Actin
Ins - - - - - + + + + + +


b

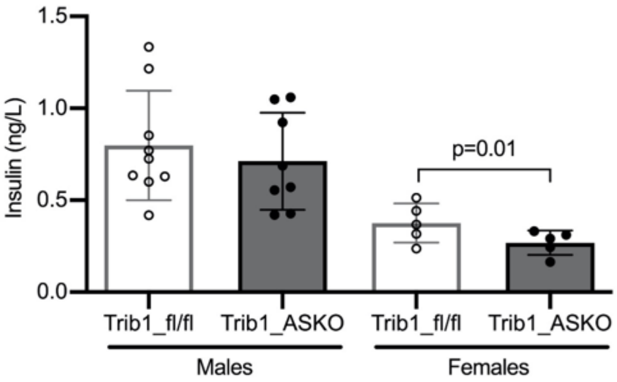

Insulin (ng/L)
1.5
1.0
0.5
0.0
p=0.01
Trib1_fl/fl
Trib1_ASKO
Trib1_fl/fl
Trib1_ASKO
Males
Females

**Figure S5**

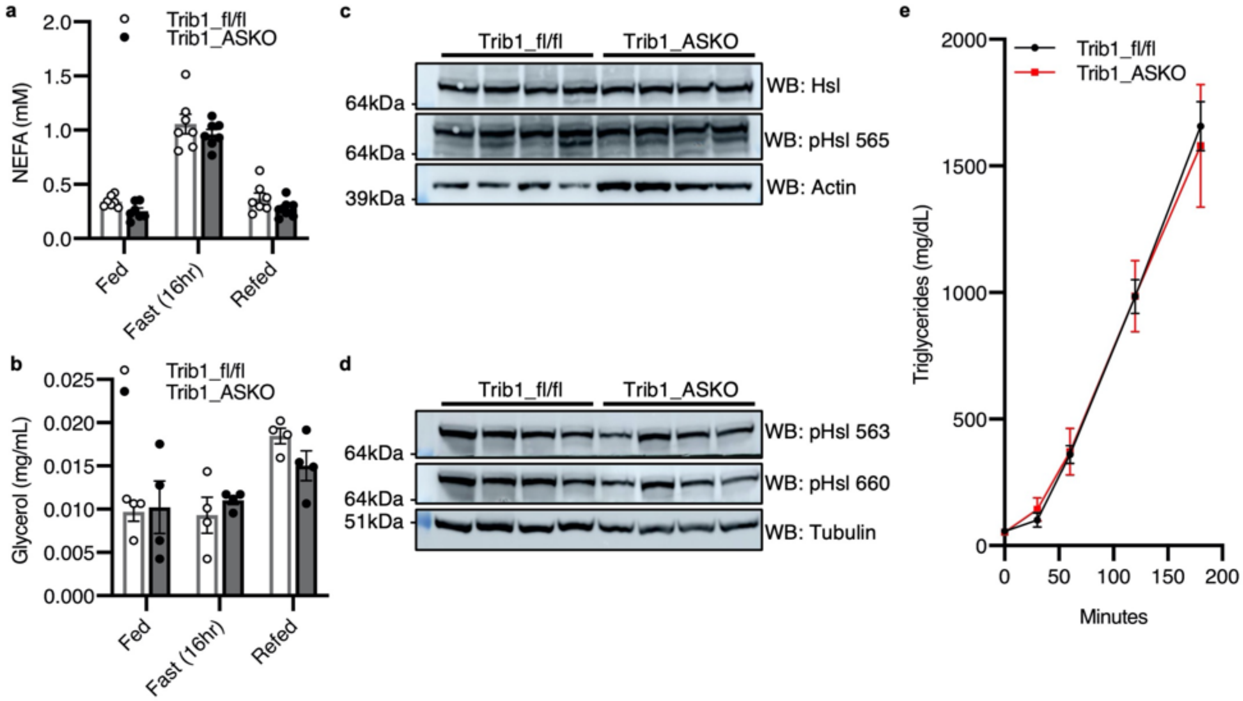
a
Trib1_fl/fl
Trib1_ASKO
NEFA (mM)
2.0
1.5
1.0
0.5
0.0
Fed
Fast (16hr)
Refed
b
Trib1_fl/fl
Trib1_ASKO
Glycerol (mg/mL)
0.025
0.020
0.015
0.010
0.005
0.000
Fed
Fast (16hr)
Refed
c
Trib1_fl/fl
Trib1_ASKO
64kDa
WB: Hsl
64kDa
WB: pHsl 565
39kDa
WB: Actin
d
Trib1_fl/fl
Trib1_ASKO
64kDa
WB: pHsl 563
64kDa
WB: pHsl 660
51kDa
WB: Tubulin
e
Trib1_fl/fl
Trib1_ASKO
Triglycerides (mg/dL)
2000
1500
1000
500
0
0
50
100
150
200
Minutes

**Table S1**

| Gene | WTMean | KOMean | log2Fold Change | FDR |
|---|---|---|---|---|
| Zfp949 | 289.63 | 972.79 | 1.75 | 3.49E-31 |
| Elovl6 | 1765.49998 | 3577.595502 | 1.017924464 | 6.19E-07 |
| Zfp9 | 70.6852799 | 187.5702233 | 1.392655502 | 0.001037606 |
| Wnk4 | 206.7594684 | 428.5849381 | 1.053514912 | 0.002585755 |
| Cyp4a31 | 828.6202647 | 1659.370849 | 1.002657112 | 0.002628167 |
| Hspa1b | 113.0580057 | 261.0773709 | 1.198030006 | 0.005313458 |
| Agap2 | 60.62624872 | 136.9006225 | 1.166696555 | 0.015898375 |
| Cgref1 | 35.78926473 | 89.0087298 | 1.314000967 | 0.031369612 |
| Ces4a | 58.03381149 | 147.8366116 | 1.33740533 | 0.043842131 |
| Adora1 | 961.420189 | 429.5204942 | -1.164833527 | 2.38E-06 |
| 9130409I23Rik | 821.1650234 | 309.1446735 | -1.407373396 | 3.40E-05 |
| Robo1 | 379.7997374 | 138.753851 | -1.455541003 | 5.21E-05 |
| Cd36 | 1375.341511 | 564.1716741 | -1.2858317 | 7.76E-05 |
| Cyp2a5 | 58151.16071 | 27733.77403 | -1.068175216 | 0.000384612 |
| Lrtm1 | 326.1802699 | 100.9248075 | -1.689193178 | 0.001255769 |
| Gsta2 | 1758.605107 | 821.0776947 | -1.100135209 | 0.002400712 |
| Gsta1 | 114.6736737 | 28.48948624 | -2.023325767 | 0.004780889 |
| Atp6v0d2 | 201.3493239 | 76.89542725 | -1.383516023 | 0.00713164 |
| Gm12718 | 166.7824469 | 64.36195564 | -1.37756025 | 0.027856712 |

**Legends:**

**Figure S1: SNPs in the 8q24 locus associate with plasma lipids, CAD, and adiponectin. a**, LocusZoom plot of SNPs in the *TRIB1* locus by TG association p-value from GLGC, *Nat. Genet.,* 2013. **b**, p-values for TC, TG, and Adpn association for select lead SNPs. Data from [a]GLGC, *Nat. Genet.,* 2013*,* [b]Van der Harst et. al*.,* Circ Res*,* 2018*,* and [c]Dastani et. al.*, Plos One*., 2012.

**Figure S2: Adipocyte-specific deletion of *Trib1* does not affect adipogenic gene expression. a-d**, Taqman qPCR for *Trib1* (**a**), *Adipoq* (**b**), *Cebpa* (**c**), and *Pparg* (**d**) in Trib1_fl/fl and Trib1_ASKO SVF at 0, 2, 5, and 7 days after differentiation induction (*n* = 2). Gene expression is expressed relative to the 0 day Trib1_fl/fl group, and depicted as mean ± s.e.m. **e**,**f**,

Representative Oil red O staining of Trib1_fl/fl (**e**) and Trib1_ASKO (**f**) SVF-derived adipocytes at Day 7 after differentiation initiation. Bar = 100 μm. Significance in all panels determined by 2-way ANOVA (Sidak's multiple comparisons test).

**Figure S3: *Trib1* ASKO adipose and adipocytes do not have decreased inflammatory gene expression.** **a**,**b**,Taqman qPCR for *Tnfa, Il6, Nos2, and Il10* in scWAT (**a**) of chow-fed Trib1_fl/fl and Trib1_ASKO male mice (n = 7) and VAT (**b**) of 12 week HFD-fed Trib1_fl/fl and Trib1_ASKO male mice (n = 7). **c**, Taqman qPCR for *Tnfa, Il1b, Il6,* and *Nos2* in Trib1_fl/fl and Trib1_ASKO SVF-derived adipocytes treated with 25 ng/mL TNFa (n=3) for 45 min. Gene expression is depicted as mean ± s.e.m. Significance in all panels determined by 2-way ANOVA (Sidak's multiple comparison test in **a**,**b** and Tukey's multiple comparison test in **c**). ns=not significant.

**Figure S4: Exploration of insulin signaling in Trib1_ASKO mice.** **a**, Immunoblot of total and phosphorylated Akt in Trib1_fl/fl and Trib1_ASKO SVF-derived adipocytes treated with 100nM insulin for 2 min. **b**, Fasting plasma insulin in HFD-fed Trib1_fl/fl and Trib1_ASKO males ($n$ = 8) and females ($n$ = 5). Data depicted as mean ± s.d. Significance determined by Student's *t* test.

**Figure S5: Markers of Lipolysis are unchanged in Trib1_ASKO adipose.** **a**, Plasma NEFA after 16 hr fast and 3 hr refeeding in Trib1_fl/fl and Trib1_ASKO mice ($n$ = 7). **b**, Plasma glycerol after 16 hr fast and 3 hr refeeding in Trib1_fl/fl and Trib1_ASKO mice ($n$ = 4). **c**, Western blot of total and phosphorylated HSL (p565) in scWAT from Trib1_fl/fl and Trib1_ASKO mice after a 4 hr fast. **d**, Western blot of phosphorylated HSL (p563 and p633) in scWAT from Trib1_fl/fl and Trib1_ASKO mice after a 4 hr fast. Blots were stripped for 15 min between consecutive staining for different phosphorylation sites. **e**, Triglyceride secretion in Trib1_fl/fl and Trib1_ASKO mice ($n$ = 6). Plasma triglycerides were measured in 4 hr-fasted mice at various timepoints after pluronic injection. Data depicted as mean ± s.d. Significance in all panels determined by Student's *t* test.

**Table S1: Differentially expressed genes in Trib1_ASKO vs. Trib1_fl/fl livers.** DESeq2 analysis of RNA-seq data from Trib1_fl/fl and Trib1_ASKO mouse hepatocytes (*n* = 4) reveals 19 differentially expressed genes.
